# EB3-informed dynamics of the microtubule stabilizing cap during stalled growth

**DOI:** 10.1101/2021.12.07.471417

**Authors:** Maurits Kok, Florian Huber, Svenja-Marei Kalisch, Marileen Dogterom

**Author notes:** Correspondence to: Marileen Dogterom.

## Abstract

Microtubule stability is known to be governed by a stabilizing GTP/GDP-Pi cap, but the exact relation between growth velocity, GTP hydrolysis and catastrophes remains unclear. We investigate the dynamics of the stabilizing cap through *in vitro* reconstitution of microtubule dynamics in contact with micro-fabricated barriers, using the plus-end binding protein GFP-EB3 as a marker for the nucleotide state of the tip. The interaction of growing microtubules with steric objects is known to slow down microtubule growth and accelerate catastrophes. We show that the lifetime distributions of stalled microtubules, as well as the corresponding lifetime distributions of freely growing microtubules, can be fully described with a simple phenomenological 1D model based on noisy microtubule growth and a single EB3-dependent hydrolysis rate. This same model is furthermore capable of explaining both the previously reported mild catastrophe dependence on microtubule growth rates and the catastrophe statistics during tubulin washout experiments.

## INTRODUCTION

Microtubules are hollow cylindrical polymers consisting of αβ-tubulin dimers arranged in a head-to-tail fashion to form protofilaments, ~13 of which typically constitute the microtubule lattice (Debs et al., 2020; Tilney et al., 1973; Zhang et al., 2015). Individual microtubules constantly switch between phases of growth and shrinkage, a fundamental process known as dynamic instability (Mitchison and Kirschner, 1984). As a major constituent of the eukaryotic cytoskeleton, microtubules are involved in many essential processes within the cell, including intracellular transport, cell division, and cell morphology (Akhmanova and Steinmetz, 2015). During these processes, dynamic microtubule ends interact with other cellular components, either through intermediary protein complexes or through direct physical contact (Colin et al., 2018; Dogterom and Koenderink, 2019; Gurel et al., 2014; Nguyen-Ngoc et al., 2007; Preciado Lopez et al., 2014; Waterman-Storer et al., 1995). Such contacts typically affect the dynamic behaviour of microtubule ends which is integral to the biological function of these interactions (Bouchet et al., 2016; Brangwynne et al., 2006; Gregoretti et al., 2006; Komarova et al., 2002; Letort et al., 2016; Tischer et al., 2009; Vleugel et al., 2016).

The biochemical mechanism behind the stochastic transition from growth to shrinkage, known as a catastrophe, is related to the progressive hydrolysis of GTP bound to β-tubulin (Carlier and Pantaloni, 1982; Nogales, 1999). During polymerization, the microtubule tip is highly dynamic due to continuous addition and removal of tubulin dimers (Gardner et al., 2011a; Kerssemakers et al., 2006; Rickman et al., 2017; Schek et al., 2007). After GTP-bound tubulin is incorporated at the microtubule tip, hydrolysis of the nucleotide followed by Pi release is hypothesized to lead to a destabilization of the lattice by a compaction around the exchangeable nucleotide (Alushin et al., 2014; Zhang et al., 2015). The delay between tubulin incorporation and hydrolysis results in a GTP/GDP-Pi enriched region at the microtubule tip, which gives rise to what is known as the stabilizing cap (Carlier and Pantaloni, 1981). Upon the loss of the stabilizing cap, a catastrophe follows upon which the strain build-up in the lattice is released during depolymerization. However, whereas this basic description of the biochemistry behind dynamic instability is generally accepted, the exact relation between GTP hydrolysis, the size of the stabilizing cap, the details of microtubule growth and the statistics of catastrophes is still not fully understood, despite the availability of a wealth of quantitative data on catastrophe statistics under different conditions (Brouhard and Rice, 2018; Ohi and Zanic, 2016).

Various estimates of the stabilizing cap size have been reported, from short caps of a few terminal tubulin layers (Brun et al., 2009; Caplow and Shanks, 1996; Drechsel and Kirschner, 1994; Karr and Purich, 1978; Walker et al., 1991) to longer caps spanning up to dozens of layers (Carlier and Pantaloni, 1981; Seetapun et al., 2012). In recent years, direct visualisation of the tubulin nucleotide state has become possible with the family of end binding proteins (EBs) that can autonomously bind to the GDP-Pi region at the microtubule tip (Maurer et al., 2011). It has been shown that the size of the EB comet at the tip correlates with the size of the stabilizing cap and consequently with microtubule stability (Duellberg et al., 2016a; Seetapun et al., 2012; Zhang et al., 2015). In fact, tubulin washout experiments using Mal3 (the EB1 homolog in yeast) suggest that not the size of the total EB binding region is decisive in preventing a catastrophe, but the presence of a critical number of unhydrolyzed subunits at the terminal ~10 tubulin layers at the microtubule tip (~130 dimers) (Duellberg et al., 2016a).

Here we use the ability of EB3 to report on the status of the stabilizing cap to investigate the detailed relation between cap dynamics and catastrophe statistics for stalling microtubules that are pushing against microfabricated barriers. A stalling microtubule exerts a pushing force that is too small to overcome its critical buckling force, resulting in the blocking of further microtubule growth. The stability of pushing microtubules has previously been studied for buckling and bending microtubules *in vitro* (Janson et al., 2003), *in vivo* (Brangwynne et al., 2007; Odde et al., 1999; Pallavicini et al., 2017), and *in silico* (Das et al., 2014; Valiyakath and Gopalakrishnan, 2018). It was established that microtubules generating pushing forces against rigid barriers *in vitro* experience an increased catastrophe frequency (Janson et al., 2003; Laan et al., 2008). This force-induced catastrophe is thought to be the result of a reduction in the addition of tubulin dimers as the microtubule growth velocity is slowed with increasing force (Janson et al., 2003; Kerssemakers et al., 2006). However, exactly how the reduction of tubulin addition in combination with nucleotide hydrolysis affects the dynamics of the stabilizing cap, and thereby determines the lifetime statistics for stalled microtubules, has remained unresolved.

We determine the lifetime statistics of stalling microtubules *in vitro* by growing microtubules against micro-fabricated barriers using GFP-EB3 as a proxy for the size of the stabilizing cap. By introducing a novel barrier design with a long overhang, we ensure that microtubule stalling can be imaged simultaneously with the EB3 signal using TIRF microscopy. We observe that microtubule stalling increases the catastrophe frequency in the absence of EB3 as previously reported (Janson et al., 2003). In the presence of EB3 the microtubule lifetime is further reduced in a concentration dependent manner. Surprisingly, the full decay of the EB3 comet during microtubule stalling does not necessarily lead to an immediate catastrophe. We compare our results to similar data obtained for freely growing microtubules under the same conditions, and then perform simulations of microtubule dynamics based on a simple model in an attempt to simultaneously explain both data sets.

Over the years, different types of models have been proposed to gain a better understanding of what triggers a catastrophe. Biochemical models rely on the hydrolysis of tubulin dimers to reduce the size of the stabilizing cap to trigger a catastrophe (Bayley et al., 1989; Brun et al., 2009; Chen and Hill, 1985; Gardner et al., 2011a; Margolin et al., 2012; Padinhateeri et al., 2012; Piedra et al., 2016; VanBuren et al., 2002). However, only with the introduction of lateral interactions between dimers in a 2D model can these models capture observed growth fluctuations (Gardner et al., 2011a) and observed microtubule lifetimes (Bowne-Anderson et al., 2013; Lee and Terentjev, 2019). Mechanochemical models additionally include the build-up of strain in the lattice and protofilament bending at the tip (Bollinger and Stevens, 2019; Coombes et al., 2013; McIntosh et al., 2018; Michaels et al., 2020; Molodtsov et al., 2005; VanBuren et al., 2005; Zakharov et al., 2015). Both types of models can explain a variety of experimental observations of dynamic instability, but they typically require many fitting parameters and do not explicitly include the highly dynamic nature of the microtubule tip. Alternatively, simple phenomenological models have been useful in obtaining an intuitive insight into the principles behind microtubule dynamics and the effect of microtubule associated proteins (Brun et al., 2009; Duellberg et al., 2016a; Flyvbjerg et al., 1996; Rickman et al., 2017).

To find a minimal model capable of explaining microtubule catastrophe statistics with the smallest possible number of fitting parameters, we use coarse-grained Monte Carlo simulations of 1D filaments (Flyvbjerg et al., 1996; Margolin et al., 2006; Padinhateeri et al., 2012; Rickman et al., 2017). We show that the lifetimes of both freely growing and stalled microtubules can be explained by a combination of random GTP hydrolysis and a parameter that characterises the noisiness of microtubule growth, a concept that was generally missing from previous phenomenological models in the description of microtubule lifetimes. We confirm that while the EB binding region is a measure for the size of the stabilizing cap, there is no one-to-one correlation between its presence and the onset of a catastrophe. Instead, the data are consistent with a catastrophe being triggered when a large enough sequence of GDP-bound tubulin dimers becomes exposed at the microtubule tip. Importantly, this 1D biochemical model can also successfully capture the previously reported catastrophe dependence on tubulin concentration, taking into account previously reported velocity-dependent growth fluctuations, and it is in good agreement with previously reported catastrophe delays after tubulin dilution. The so-called “ageing” of microtubules, referring to the observed increase of the catastrophe frequency with microtubule age (Coombes et al., 2013; Duellberg et al., 2016b; Gardner et al., 2011b; Odde et al., 1995), is not a feature of our simple model, but this behaviour naturally emerges by additionally assuming that microtubule growth fluctuations increase with microtubule age.

## RESULTS

### *In vitro* reconstitution of microtubule stalling

To investigate the stability of stalling microtubules in the presence of different concentrations of GFP-EB3, we analysed the dynamics of microtubules growing against micro-fabricated barriers using an *in vitro* reconstitution assay (Bieling et al., 2007; Kalisch et al., 2011). We assume each microtubule to consist of three regions: 1) a GTP rich terminal region where protofilament growth takes place (Maurer et al., 2014; McIntosh et al., 2018), 2) a region containing the intermediate GDP-Pi state to which EB3 preferably binds (Maurer et al., 2011), and 3) the GDP lattice (Fig. 1A). The presence of EB3 is known to increase both the GTP hydrolysis rate and the microtubule growth velocity by respectively compacting the microtubule lattice and closing the lattice seam (Maurer et al., 2014; Zhang et al., 2015). Microtubules were nucleated from GMPCPP-stabilized seeds and imaged with Total Internal Reflection Fluorescence (TIRF) microscopy. The barriers were composed of 100 nm SiO_2_ deposited on a glass coverslip with an amorphous silicon carbide (SiC) overhang, approximately 1.5 μm long, to trap the microtubules and force them to grow into the barriers (Fig. 1BC). SiC is a mechanically stable, optically transparent material (wavelengths > 0.5 μm) with similar passivation and functionalization properties as SiO_2_ due to a very thin native oxide layer on its surface (Coletti et al., 2007; Dhar et al., 2009). Although fabrication of the barriers requires a thin 10 nm layer of SiC on the glass surface (see Methods for details), microtubules can be imaged successfully with TIRF microscopy (Fig. 1D and S1). This novel barrier design enables high resolution imaging with TIRF microscopy as the microtubules are forced to remain within 100 nm from the surface during barrier contact, eliminating fluctuations perpendicular to the surface. The width between two barriers is 15 μm, chosen to keep the microtubules short and thereby reduce the probability of observing slipping and buckling events (Janson et al., 2003). All experiments were performed in the presence of 15 μM tubulin, and 0, 20, 50, or 100 nM GFP-EB3.

**Figure 1.**
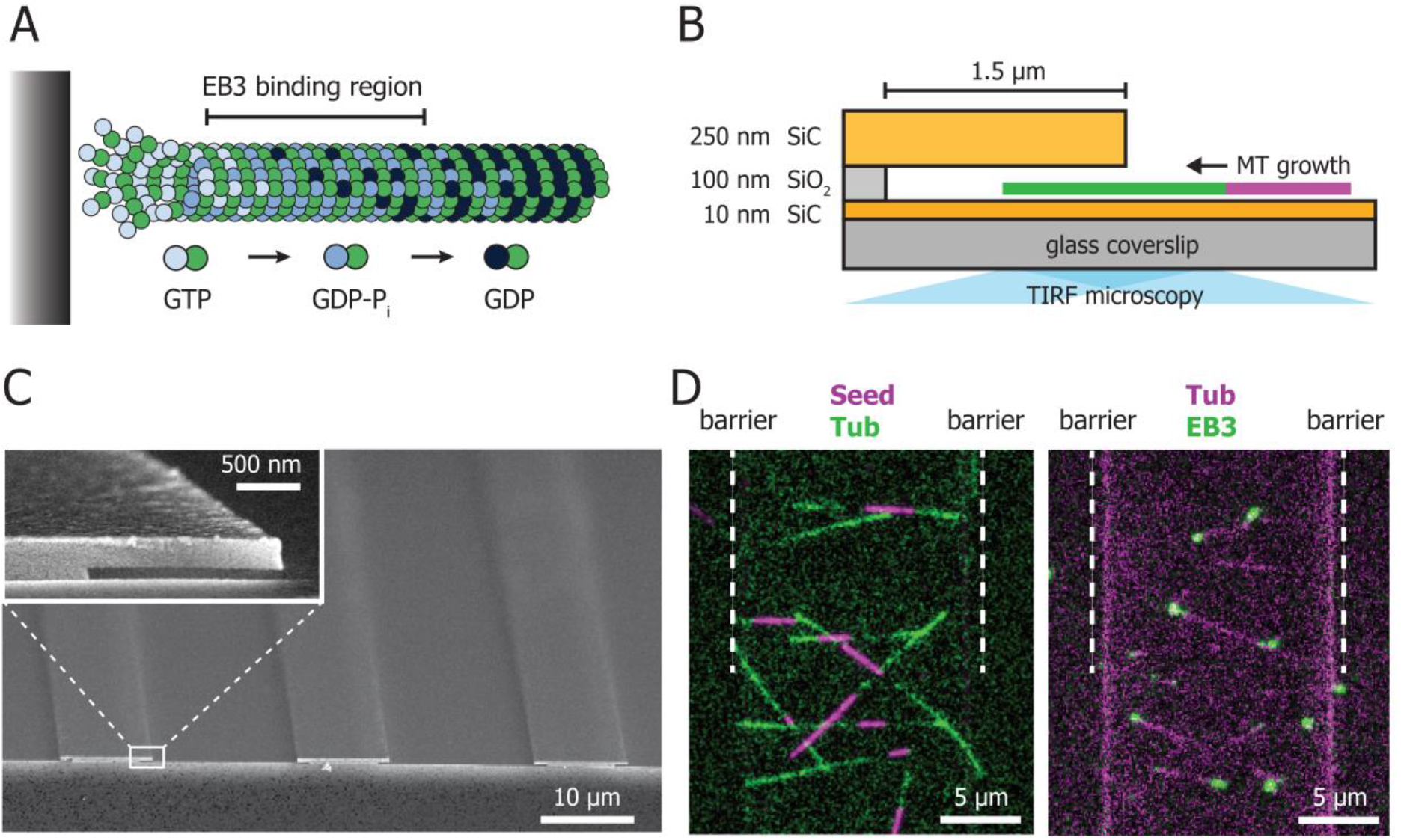
*In vitro* reconstitution of microtubule stalling. **(A)** Schematic depiction of the nucleotide distribution at the microtubule tip. Progressive hydrolysis of GTP-tubulin after incorporation leads to a destabilized lattice that is stabilized by a GTP/GDP-Pi cap. EB3 chiefly binds to the GDP-Pi rich region. During microtubule-barrier contact, reduction of microtubule growth and ongoing hydrolysis are hypothesized to lead to an accelerated loss of the protective cap. **(B)** Schematic of the micro-fabricated barrier with a 1.5 μm overhang made of silicon carbide (SiC). The barrier itself is composed of SiO_2_ and is 100 nm high, forcing the growing microtubules to remain within the TIRF illumination field. Microtubules are grown from stabilized GMPCPP seeds. **(C)** Scanning Electron Microscope image of two channels with barriers. The insert shows a zoom of the barrier with a 1.5 μm SiC overhang. **(D)** TIRF images of the micro-fabricated channel enclosed by two barriers (white dotted lines). On the left microtubules (green) are nucleated from GMPCPP-stabilized seeds (magenta) towards the barriers and on the right microtubules (magenta) polymerize towards the barriers in the presence of GFP-EB3 (green). See also Fig S1 and Videos S1 and S2.

### Complete EB3 decay is neither needed for, nor always immediately followed by a catastrophe

The microtubule-barrier contact events leading to a stalling microtubule were analysed with kymographs to obtain the contact duration and GFP-EB3 comet intensity prior and during contact (Fig. 2A and S1, see Methods for details). Any contact events leading to microtubule buckling or sliding along the barrier were excluded from the analysis. From the moment of barrier contact, the EB3 intensity at the microtubule tip decreased until the onset of catastrophe (Fig. 2B, S1D). We observed that for ~65% of all stalling events, a catastrophe occurred while the EB comet was still decaying (> 10% of the EB signal remaining) (Fig 2C, right). For those events, the mean comet intensity at the moment of catastrophe was 16% of the pre-contact mean. For the remaining ~35% of stalling events we found a decay of the EB comet to a steady near-zero value which did not immediately lead to catastrophe. Instead, the microtubules remained in contact with the barrier for some time, even in the absence of an observable EB3 comet (Fig. 2C, left). Fitting the average EB decay from the moment of barrier contact with a mono-exponential function, shows that the decay rate increases with EB3 concentration: 0.24 s^-1^, 0.29 s^-1^, and 0.40 s^-1^ for 20, 50, and 100 nM EB3 respectively (Fig. 2D). The presence of EB3 thus accelerates the decay of the EB comet during stalling events as predicted from its increasing effect on the GTP hydrolysis rate (Maurer et al., 2014; Zhang et al., 2015).

**Figure 2.**
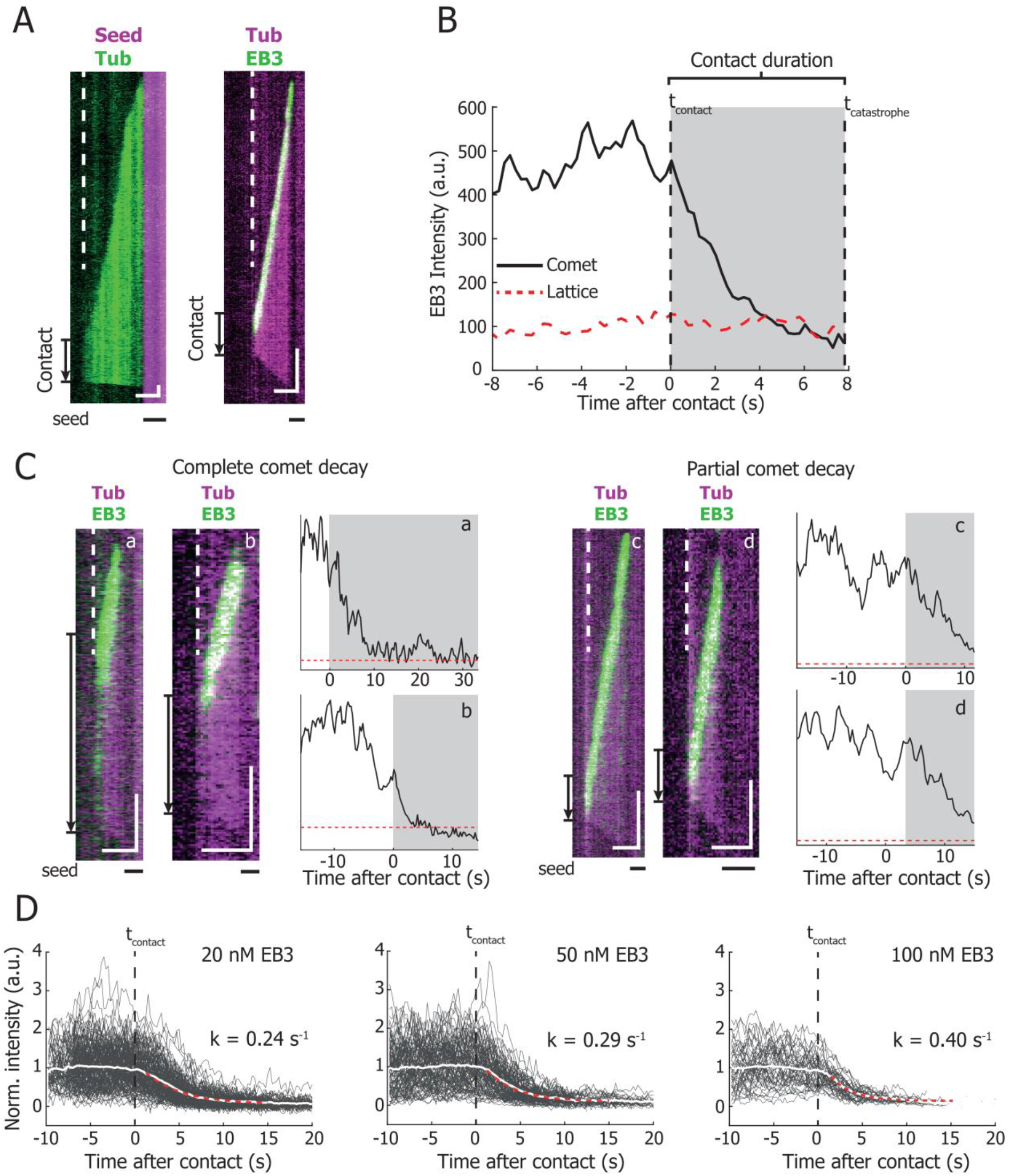
Microtubule stalling events *in vitro*. **(A)** Representative kymographs of a microtubule-barrier contact event in the presence of 15 μM Hilyte488-labelled tubulin (left) and in the presence of 15 μM rhodamine-labelled tubulin with 20 nM GFP-EB3 (right). The dotted line denotes the position of the SiO_2_ barrier. The duration of barrier contact is indicated with an arrow. Scale bars: 2 μm (horizontal) and 10 s (vertical). **(B)** Mean intensity of the EB3 comet and the EB3 signal on the microtubule lattice of the kymograph on the right in **(A)**. From the moment of microtubule-barrier contact (*t_contact_*), the EB3 comet signal decays to the level of the microtubule lattice, ultimately resulting in the onset of a catastrophe (*t_catastrophe_*) after 7.75 seconds. **(C)** Several examples of stalling microtubules with their respective comet intensity traces. Traces **a-b** show a full comet decay during barrier contact before the onset of a catastrophe, whereas the comet in traces **c-d** only partially decays. All traces were in the presence of 15 μM tubulin. Additionally, trace **a** contained 20 nM EB3, trace **b** and **c** 100 nM EB3, and trace **d** 50 nM EB3. Arrows and shaded regions illustrate the duration of microtubule stalling event. Scale bars: 2 μm (horizontal) and 10 s (vertical). **(D)** Normalized comet intensity traces of stalling microtubules in the presence of 20, 50, and 100 nM EB3, aligned on the moment of barrier contact (*t_contact_*). The mean decays were fitted with a mono-exponential model and show an increasing decay rate with increasing EB3 concentrations, resulting in decay rates of 0.24 s^-1^, 0.29 s^-1^, and 0.40 s^-1^ for 20, 50, and 100 nM respectively. Number of stalling events analysed: 20 nM, n = 151, 50 nM, n = 104, and 100 nM, n = 92.

### Monte Carlo simulation of microtubule catastrophes

To determine whether microtubule lifetime statistics of stalling microtubules can be understood solely by a single stochastic hydrolysis step combined with net stalling of noisy microtubule growth, we performed minimalistic Monte Carlo simulations of both free and stalled microtubule growth. Microtubules were treated as 1D filaments with subunits of 8/13 nm comprising two distinct states: GTP/GDP-Pi and GDP (Fig. 3A). We decided to ignore the initial transition from GTP to GDP-Pi, which was reported to be much faster than the GDP-Pi to GDP transition (Kim and Rice, 2019; Maurer et al., 2014; Rickman et al., 2017). We saw this justified by the fact that our key observations changed only very moderately when the first transition was included explicitly, whereas omitting this transition reduced the number of model parameters. Simulated microtubules grow by addition of GTP/GDP-Pi subunits which subsequently undergo random hydrolysis to GDP with rate *k_hyd_* (Fig. 3A). We treat microtubule growth as a discrete, biased Gaussian random walk (Antal et al., 2007; Flyvbjerg et al., 1996), inspired by experimental observations that revealed a substantial diffusive character of the growing microtubule tip (Gardner et al., 2011a; Kerssemakers et al., 2006; Rickman et al., 2017; Schek et al., 2007). Following this model, tip growth is fully characterized by the experimentally measured mean growth velocity 〈*V*〉 and the diffusion constant *D_tip_*, which may also result in occasional negative growth excursions (Fig. 3B). A catastrophe is triggered when a stabilizing cap (*L_cap_*) that results from remaining GTP/GDP-Pi subunits is lost due to a negative growth excursion and/or random hydrolysis. We assume that the nucleotide state of the tubulins at the very tip of the microtubule are the most relevant for stability: a catastrophe is triggered when the number of uninterrupted GDP subunits at the very tip of the microtubule is equal or greater than *N_unstable_* (Brun et al., 2009; Padinhateeri et al., 2012), independent of how many GTP/GDP-Pi subunits remain elsewhere in the lattice (Fig 3CD). The length of the stabilizing cap is thus determined by the distance between the position of the microtubule tip and the position along the lattice where for the first time an uninterrupted sequence (or ‘island’) of GDP units equal or greater than the fitting parameter *N_unstable_* is found (Fig. 3C). Depolymerization and rescues are not considered in the model. To exclude nucleation kinetics from the simulated lifetimes, a microtubule is considered to grow after reaching a length of 250 nm. The simulation then only requires the three fitting parameters *k_hyd_*, *D_tip_*, and *N_unstable_*, all of which can be verified with experimental data (see below). The experimental EB3 intensity at the microtubule tip can be compared to the total number of GTP/GDP-Pi-state subunits in the simulated microtubules (Fig. 3DE, bottom).

**Figure 3.**
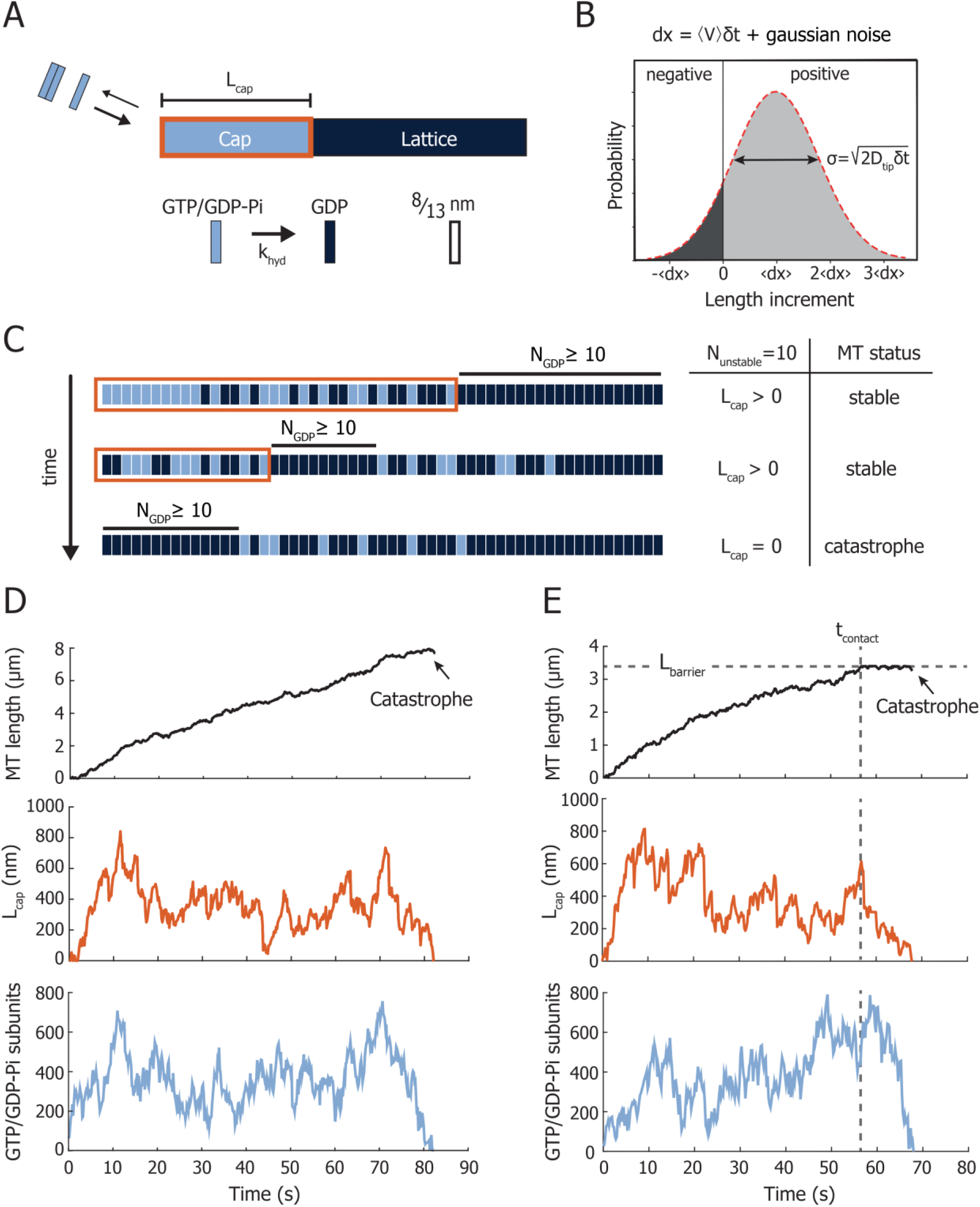
Monte-Carlo simulation of microtubule dynamics. **(A)** The microtubule is simulated as a one-dimensional lattice with two states, a GTP/GDP-Pi and a GDP state, that are approximately distributed in a cap with length *L_cap_* and a lattice region respectively. The size of each subunit is 8/13 nm. Uncoupled stochastic hydrolysis matures the GTP/GDP-Pi-state into the GDP-state with rate *k_hyd_*. **(B)** Microtubule growth is determined by a mean growth velocity 〈*V*〉 with added Gaussian noise characterized by *D_tip_*, resulting in stochastic tip elongation following a biased random walk. The diffusive character of the tip can also produce negative growth excursions. **(C)** Detailed schematic of the nucleotide composition of the microtubule tip. The size of the stabilizing cap (orange) is defined as the region between the microtubule tip and the position along the lattice where for the first time an uninterrupted sequence of GDP subunits is equal or greater than *N_unstable_*. A catastrophe is triggered when the number of uninterrupted GDP subunits at the very tip of the microtubule is equal or greater than *N_unstable_*. As example, the case for *N_unstable_* = 10 is shown. **(D)** Simulated microtubule growing event. **(top)** During the noisy microtubule growth, the microtubule length follows a biased random walk. **(middle)** When the size of the stabilizing cap (*L_cap_*) is reduced to zero, i.e. when the number of uninterrupted terminal GDP subunits is equal or greater than *N_unstable_*, a catastrophe follows, and the simulation is terminated. **(bottom)** Total number of GTP/GDP-Pi subunits in the simulated microtubule, which can be compared with the experimentally obtained EB3 signal. **(E)** Simulated microtubule stalling event. Figures show the simulated microtubule length **(top)**, the size of the stabilizing cap (*L_cap_*) **(middle)**, and the total number of GTP/GDP-Pi subunits in the simulated microtubule **(bottom)**. Barrier contact is simulated by restricting the maximum length of the microtubule to *L_barrier_*. As the microtubule can undergo occasional negative growth excursions due to noisy growth, the microtubule length can still fluctuate during barrier contact.

Microtubule stalling is simulated by introducing a fixed maximum length *L_barrier_* (Fig. 3E). Any growth excursions that would bring the microtubule length to *L* > *L_barrier_* are truncated to this maximum length. Since tip fluctuations also include occasional negative growth excursions (Fig. 3B), fluctuations of the tip position continue after barrier contact (Fig. 3E).

### Obtaining the simulation parameters

Our 1D model relies on three fitting parameters, *D_tip_, k_hyd_*, and *N_unstable_*. Based on existing literature (Maurer et al., 2014), we expected that adding EB3 would have an effect on the transition from GTP/GDP-Pi to GDP. Since EB3 also affects the microtubule growth velocity, we also expected that growing microtubules may display different growth fluctuations at different EB concentrations. It hence appeared reasonable to keep *k_hyd_* and *D_tip_* as free fitting parameters, while keeping *N_unstable_* as a global fitting parameter that is independent of the presence of EB3. We obtained values for the mean growth velocity 〈*V*〉 of 1.7, 2.8, 2.8, and 3.7 μm/min for 0, 20, 50, and 100 nM EB3 respectively and a global mean seed-barrier distance *L_barrier_* of 3.4 μm from the experimental data.

To find good fitting values, we performed systematic parameter scans across a range of *D_tip_* and *k_hyd_*, simulating 500 microtubule growth events for each parameter combination. We simulated both freely growing and stalling microtubules and compared the distributions with the respective experimental distributions. Using a Kolmogorov-Smirnov test as a measure of the similarity between the simulated and experimental distributions, we obtained heatmaps with (normalized) similarity parameters for each compared distribution (Fig. 4A). We also included a comparison between the simulated GTP/GDP-Pi decay and the experimental EB decay rate during stalling (Fig. 2D). The resulting range of *k_hyd_* values that captured the experimental decay rates was used to restrict the range of possible *k_hyd_* values for the comparison of simulated and experimental lifetime distributions. The parameter set best capturing all three comparisons was then found by calculating the product between the heatmaps within the range allowed by the decay rates (Fig. 4A and S2).

**Figure 4.**
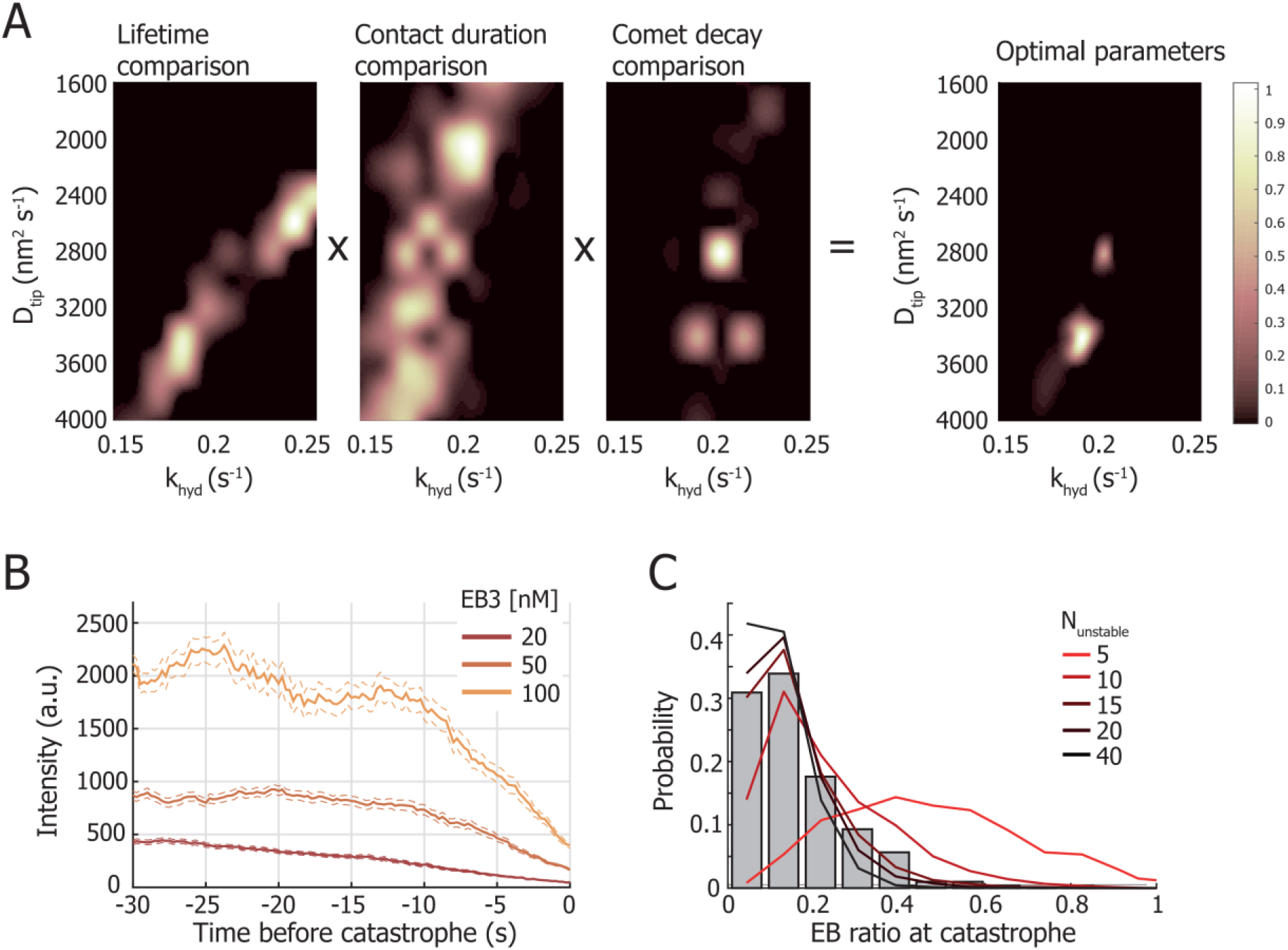
Parameter determination of the 1D model. **(A)** The diffusion constant *D_tip_* and hydrolysis rate *k_hyd_* are determined by comparing the simulated microtubule lifetime distributions, the contact duration distributions, and the decay of the EB signal during contact with the respective experimental distributions. The lifetime and contact duration distributions are compared with the experimental distributions using a Kolmogorov-Smirnov test. A cubic interpolation of the similarity values is captured in (normalized) heatmaps. The comet decay rate comparison is obtained by evaluating the absolute difference between simulated rates and experimental rates. Evaluating the product of the three heatmaps results in a parameter pair of *D_tip_* and *k_hyd_* best describing the datasets. The shown heatmaps are of 20 nM EB3. See also Fig. S2. **(B)** The mean experimental EB3 signal during barrier contact, aligned on the moment of catastrophe for 20 (n=151), 50 (n=104), and 100 nM (n=92) EB3. The ratio between the steady-state EB3 signal prior to catastrophe (from −30 to −15 seconds) and at the moment of catastrophe can be compared with the simulated data. The dotted lines denote the SEM. **(C)** Histogram of the ratio between the mean comet intensity during steady-state growth and the comet intensity at the moment of catastrophe. The data is pooled from all experimental datasets of 20, 50, and 100 nM EB3 (n=347). The lines show the simulated GTP/GDP-Pi ratio for *N_unstable_* values of 5, 10, 15, 20, and 40. We find that a minimum *N_unstable_* value of 15 is required to capture the distribution of pooled experimental EB3 ratios.

### Determining the catastrophe threshold for the 1D model

To determine the catastrophe threshold governed by *N_unstable_*, we made use of the experimentally observed decay of the mean EB signal at the barrier (Fig. 2D). Our analysis yielded simultaneous fits of *D_tip_* and *k_hyd_* that were in very good agreement with our experimental observations across a wide range of *N_unstable_* values. To determine an ideal value for *N_unstable_* to match our data, we looked at the ratio between the mean EB3 signal during steady-state growth and the EB3 signal at the moment of catastrophe. This would give us a measure of what fraction of GTP/GDP-Pi subunits was on average hydrolysed at the moment a catastrophe occurred (Fig. 4B). Higher values for *N_unstable_* gave rise to a longer stabilizing cap, resulting in a higher ratio of hydrolysed subunits in the cap at the moment of catastrophe. Comparing the combined distributions of the EB3 ratios with simulated ratios for several *N_unstable_* values results in a minimum *N_unstable_* value of ~15 subunits (Fig. 4C).

### The 1D model can successfully capture microtubule lifetimes

Figures 5A and 5B show the experimental cumulative fraction of the lifetimes of freely growing microtubules and of the stalling duration respectively (bold lines), in the presence of 15 μM tubulin and 0, 20, 50, and 100 nM GFP-EB3. The distribution of free lifetimes was determined using microtubules growing parallel to the barriers. By bootstrapping each simulated distribution obtained with the best-fitting parameters (Fig. 5C), we show 25 simulated traces containing an equal number of data points as the experimental dataset (thin lines). The variability in the simulated distributions provides a good visual reference of the similarity between experiment and simulation (Fig. 5AB and S2). The distributions show that an increasing concentration of EB3 decreases the contact duration (Fig. 5BC). In the absence of EB3 the contact duration is 30.8 ± 1.3 seconds (median ± SE), whereas in the presence of 20, 50, and 100 nM GFP-EB3 the contact duration is reduced to respectively 13.0 ± 0.7, 9.1 ± 0.8, and 4.1 ± 0.3 seconds (median ± SE). The simulated distributions capture the data well and show that free microtubule lifetimes and microtubule stalling can indeed be simultaneously captured with a 1D model comprising three parameters (Fig 5A-C). From the fits, we find that with increasing EB3 concentration, *k_hyd_* increases and *D_tip_* decreases (Fig. 5C).

**Figure 5.**
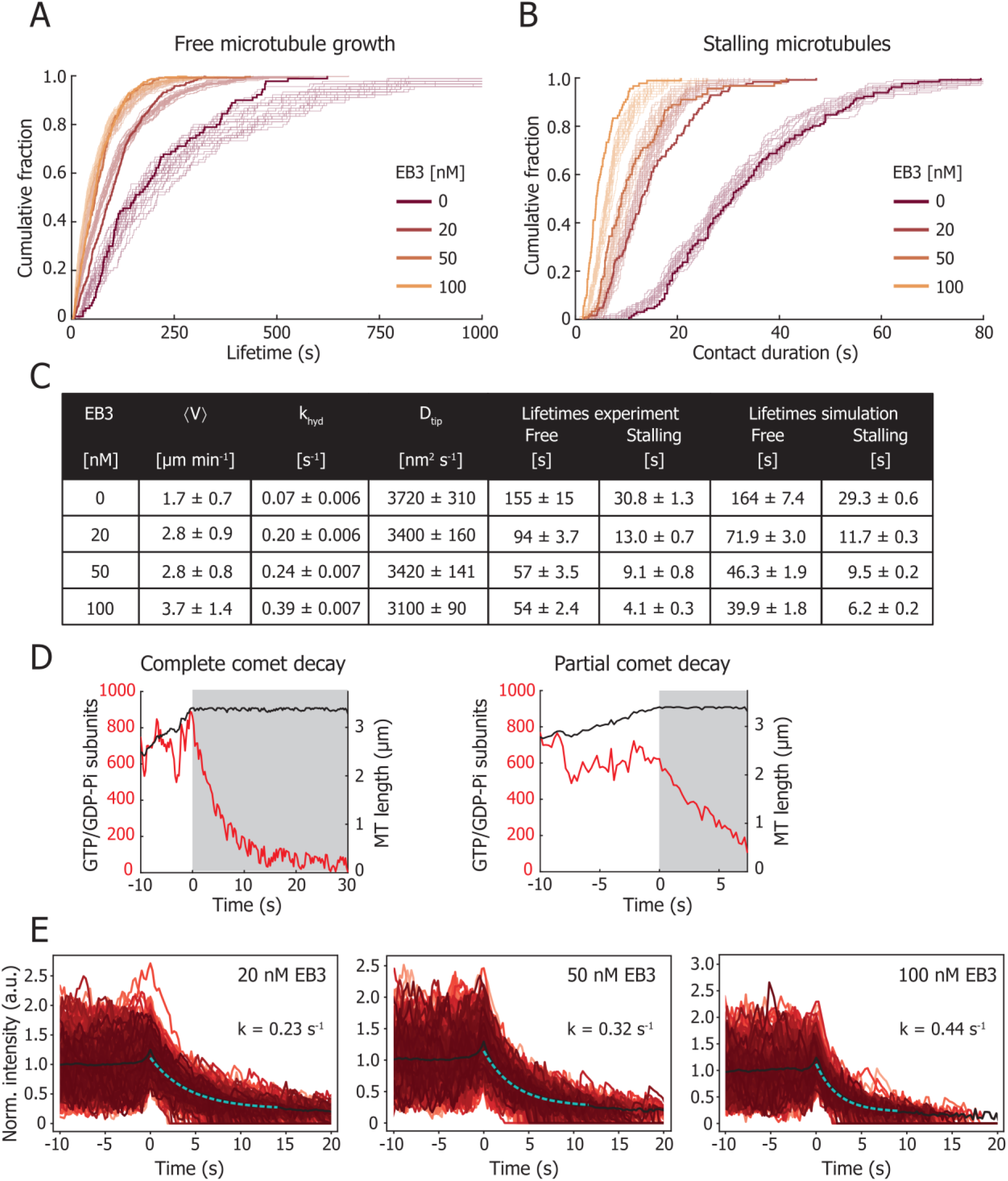
Free microtubule growth and microtubule stalling. **(A)** The cumulative fraction of the lifetimes of freely growing microtubules at increasing concentrations of 0 nM (n=90), 20 nM (n=384), 50 nM (n=262), and 100 nM (n=398) EB3. All data were obtained with 15 μM tubulin. The bold lines show the experimental data and the thin lines show 25 bootstrapped simulated distributions of equal number of datapoints as the experimental distribution. **(B)** The cumulative fraction of the microtubule-barrier contact durations at increasing concentrations of 0 nM (n=131), 20 nM (n=126), 50 nM (n=90), and 100 nM (n=90) GFP-EB3. All data were obtained with 15 μM tubulin. The bold lines show the experimental data and the thin lines show 25 bootstrapped simulated distributions of equal number of datapoints as the experimental distribution. **(C)** Table with the growth velocity 〈*V*〉 (mean ± std), hydrolysis rate *k_hyd_* (mean ± 95% CI), and diffusion constant *D_tip_* (mean ± 95% CI) as determined to simulate the lifetimes of freely growing microtubules and the contact duration of stalling microtubules at each EB concentration (median ± SE). **(D)** Examples of simulated stalling events, showing the microtubule length and the total number of GTP/GDP-Pi subunits in the lattice. The left trace shows a full comet decay during barrier contact before the onset of a catastrophe, whereas the comet in traces on the right only partially decays. **(E)** Simulated decay of GTP/GDP-Pi subunits during microtubule stalling at increasing EB concentration. Each dataset contains 1000 simulated events.

Additionally, we find both fully and partially decayed GTP/GDP-Pi intensities at the moment of catastrophe, in agreement with experimentally observed event types (Fig. 5D and 2C). Agreement between the experimental and simulated distributions of the remaining EB3 signal at the moment of catastrophe was ensured with *N_unstable_* = 15 (Fig. 4C). The mean decay rate of GTP/GDP-Pi subunits during stalling also matches the experimental dataset well and increases with increasing EB3 (Fig. 5E and 2D). The simulated barrier contact events furthermore show a similar noisy comet intensity before catastrophe, confirming that the size of the microtubule stabilizing cap fluctuates with time.

### Simulation of tubulin washout

Recent experiments using microfluidics assisted washout of tubulin *in vitro* have shown that a minimal stable cap has a length of ~10 tubulin layers at most, of which 15-30% dimers remain unhydrolysed (Duellberg et al., 2016a). The observed delay between tubulin washout and microtubule catastrophe is reported to be ~7 seconds (Walker et al., 1991) and shown to depend on the pre-washout growth velocity (Duellberg et al., 2016a). To verify the ability of our model to describe tubulin washout experiments, we simulated tubulin washout with our obtained parameter set (Fig. 5C). To simulate washout, we prohibit any growth of the microtubule tip after 40 seconds, but still allow microtubules to undergo negative growth excursions (Fig 3B). To compare our results to published washout parameters (20 μM tubulin with 0 and 200 nM of Mal3) (Duellberg et al., 2016a), we simulate tubulin washout for 15 μM tubulin in the presence of 0 and 20 nM EB3, resulting in a comparable growth velocity and hydrolysis rate. The difference between the concentration of Mal3 (fission yeast homolog of EB1) and EB3 required to obtain a similar hydrolysis rate can be explained by the intrinsic structural differences (Roth et al., 2018; von Loeffelholz et al., 2017). Our simulation of tubulin washout showed a delay between washout and catastrophe of 10.2 ± 3.2 and 5.5 ± 1.8 seconds (mean ± std) for 0 and 20 nM EB3 (Fig. 6A), similar to the reported values of 7.3 and 3.5 seconds for 0 and 200 nM Mal3 (Duellberg et al., 2016a). In addition, it was reported that microtubule growth is not simply paused after washout, but that microtubules slowly shrank prior to catastrophe. During the simulated washout delay, we measured a slow decrease in microtubule length of 253 ± 71 nm (mean ± std) for 20 nM EB3 (Fig. 6B), comparable to the reported 165 ± 105 nm for 200 nM Mal3 (Duellberg et al., 2016a). Furthermore, we find that the simulated decay rate of 0.30 s^-1^ (20 nM EB3) of GTP/GDP-Pi subunits from the moment of tubulin washout is in agreement with the reported 0.33 s^-1^ (200 nM Mal3) (Fig. 6C).

**Figure 6.**
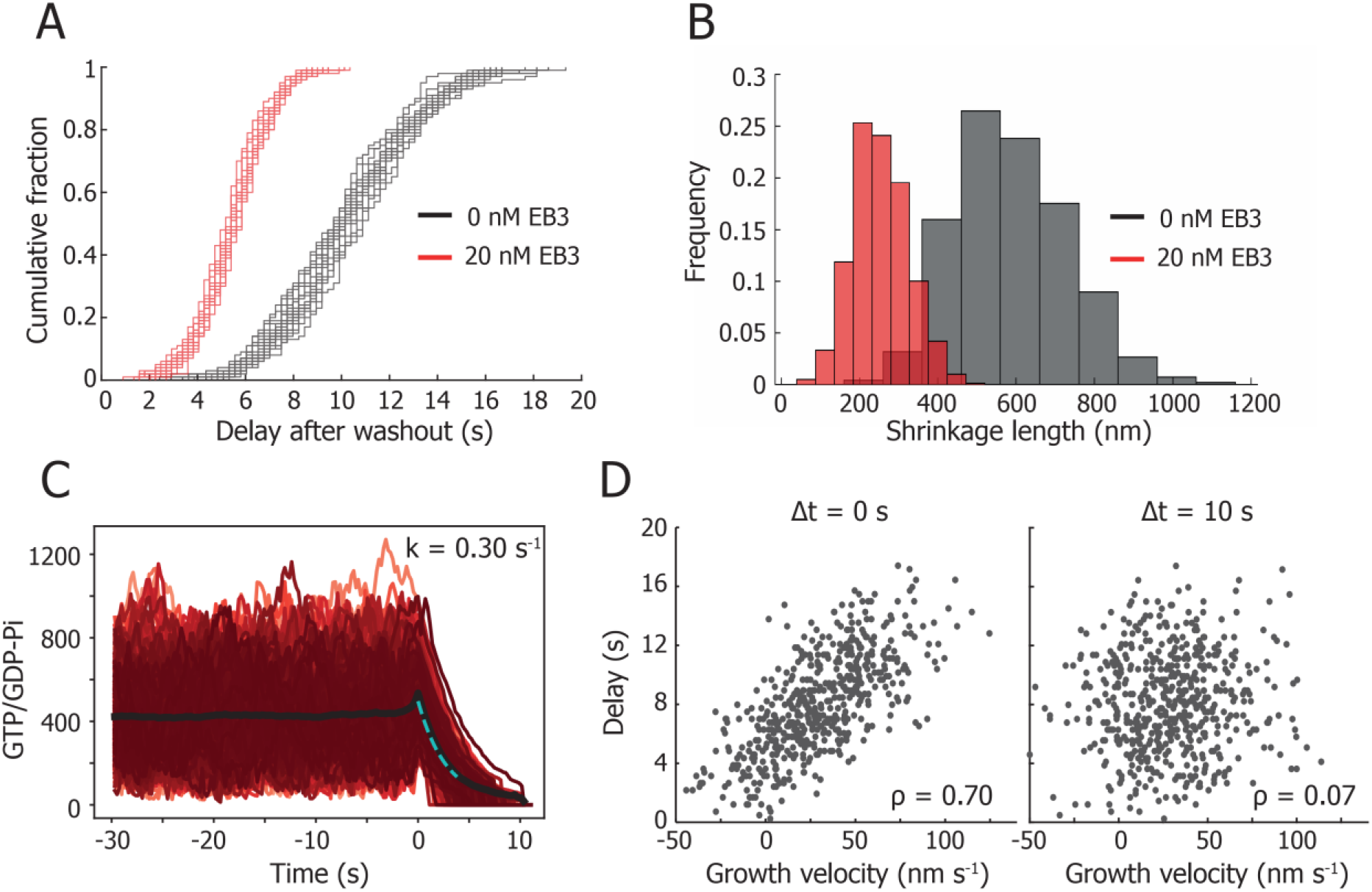
Simulating tubulin washout. **(A)** Simulation of tubulin washout following the parameters in Fig. 5C for 0 and 20 nM EB3. For both conditions, 25 bootstrapped distributions of 100 data points are shown. The mean delay duration between washout and catastrophe is 10.3 ± 3.2 and 5.5 ± 1.8 seconds for 0 and 20 nM EB3 respectively (mean ± std). **(B)** Simulation of tubulin washout following the parameters in Fig. 5C for 0 and 20 nM EB3. The mean shrinkage length of the microtubule between washout and catastrophe is 588 ± 144 and 253 ± 71 nm respectively (mean ± std). **(C)** Simulation of the number of GTP/GDP-Pi subunits before and during tubulin washout following the parameters in Fig. 5C for 20 nM EB3. Fitting the loss of GTP/GDP-Pi subunits from the moment of washout gives a decay rate of 0.30 s^-1^. **(D)** Scatter plot of the simulated delay time dependency on the growth velocity before tubulin washout. Growth velocities measured immediately before tubulin washout (Δ*t* = 0 seconds) show a strong correlation, which is lost for growth velocities measured Δ*t* = 10 seconds before washout. The mean growth velocity is calculated from a 10 second time window. ρ is Spearman’s rank correlation coefficient.

Our simulated data also captures the reported positive correlation between microtubule stability and growth velocity (Fig. 6D) (Duellberg et al., 2016a). The simulated washout delay increases with an increasing growth velocity as measured immediately prior to washout (Spearman correlation coefficient of *ρ* = 0.70). However, the correlation is lost when the growth velocity is measured 10 seconds before washout (*ρ* = 0.07), in agreement with published results (Duellberg et al., 2016a). We conclude that our model is thus capable of accurately capturing tubulin washout experiments.

### The 1D model successfully captures the mild catastrophe dependence on microtubule growth rates

As an additional verification of our model, we simulated microtubule lifetimes and stalling durations based on our previous experimental data (Fig. 7) (Janson et al., 2003). Our model can simultaneously capture the reported mild reduction in catastrophe rate with increasing growth velocity as well as the distribution of stalling durations (Fig. 7AB). To explain these data, we had to assume a velocity-dependent tip noise (at a fixed *k_hyd_* of 0.10 s^-1^), which is in line with the reported dependence of the tip noise on the growth velocity (Fig. 7C) (Gardner et al., 2011a; Rickman et al., 2017). This relationship shows a linear dependence of the tip noise on the growth velocity, as has been developed for a 1D model (Gardner et al., 2011a; Rickman et al., 2017):

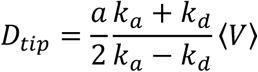

where *a* is the size of a dimer, *k_a_* the tubulin addition rate, and *k_d_* the tubulin dissociation rate. We thus conclude that our 1D model can describe the reported mild dependence of the microtubule lifetimes on the growth velocity.

**Figure 7.**
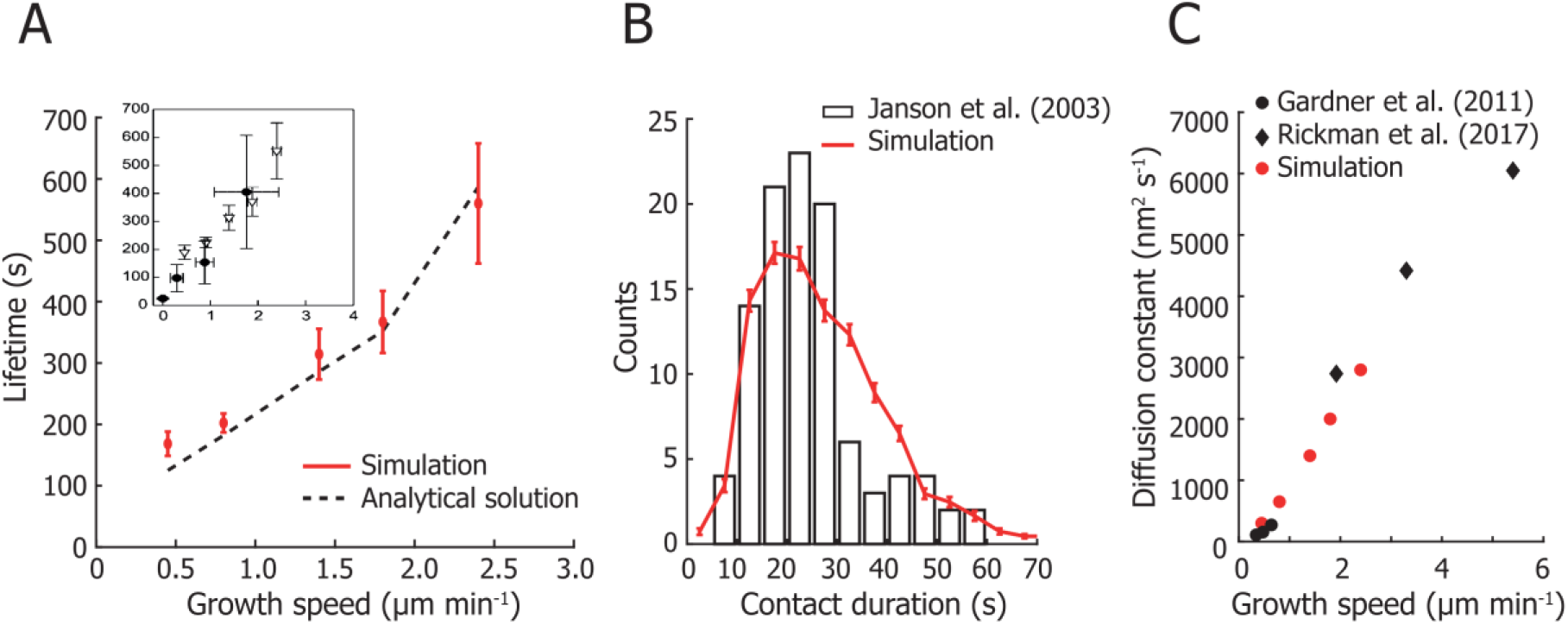
Model evaluation. **(A)** Simulation of microtubule lifetimes for increasing growth velocities. The simulation growth velocities were obtained from (Janson et al., 2003) and combined with a global value for *k_hyd_* of 0.10 s^-1^ and the determined *N_unstable_* of 15. The insert shows the experimental lifetimes for freely growing microtubules (triangles) and for buckling microtubules (dots) (Janson et al., 2003). Our simulation gives a similarly mild suppression of catastrophes with increasing tubulin concentrations, with the growth velocities of 0.45, 0.8, 1.4, 1.8, and 2.4 μm min^-1^ corresponding to tubulin concentrations of 7.2 (n=58), 10 (n=152), 15.2 (n=49), 20 (n=51), and 28 μM (n=30). Simulated lifetimes are given as mean ± SEM with the same number of datapoints as the experimental values. The mild catastrophe suppression is also captured by the analytical solution of our model (see Methods). **(B)** Histogram of the pooled stalling duration with 103 events measured at 15.2, 20, and 28 μM from (Janson et al., 2003) and the simulated stalling duration. The simulated values represent mean ± std for n = 103 events. **(C)** The diffusion constant of the microtubule tip required to simulate the microtubule lifetimes in **(A)** based on data from (Janson et al., 2003). These values agree with reported values by (Gardner et al., 2011a; Rickman et al., 2017) and follow a linear dependency on the growth speed.

## DISCUSSION

### EB3 enhances catastrophes for stalling microtubules

Using novel micro-fabricated barriers in conjunction with TIRF microscopy, we studied the duration of barrier contact as well as the dynamics of the EB3 comet during microtubule stalling. We confirm that stalled microtubules undergo a catastrophe after 30.8 ± 1.3 seconds (median ± SE) in the absence of EB3, comparable to previously measured values (Fig. 7B) (Janson et al., 2003). The presence of EB3 further enhances catastrophes in a concentration dependent manner which results in up to five times shorter microtubule contact times at the barriers (Fig 5A-C). In earlier unpublished experiments, we made similar observations for stalling microtubules in the presence of Mal3 (Fig. S4). These shorter contact times are accompanied by an increase in the decay rate of the EB3 comet (Fig. 2D).

Additionally, we developed a simple phenomenological computational model that predicts catastrophe statistics based on parameters related to random (uncoupled) GTP hydrolysis and fluctuations in microtubule growth. Fitting the model to the data suggests that the size of the growth fluctuations (*D_tip_*) decreases in the presence of EB3 (Fig. 5C). This effect would support the hypothesis that the increase in growth velocity due to the presence of EB3 is the result of a lower tubulin dissociation rate at the microtubule tip. If we assume that the tubulin association rate at the microtubule tip only depends on the soluble tubulin concentration and is therefore not affected by EB3, we would indeed expect the resulting tip noise *D_tip_* to be smaller with increasing concentrations of EB3. This effect could originate from EB3 binding in between protofilaments and reducing tip fluctuations or from the hypothesis that EB3 increases the growth velocity by closing the lattice seam (Zhang et al., 2015).

### A 1D phenomenological model successfully describes microtubule lifetimes, stalling, and tubulin washout

We developed a simple phenomenological computational model that can capture a very rich set of experimental data on dynamic microtubules. Its sole dependence on (velocity-dependent) tip noise and random hydrolysis makes it possible to build an intuition of key processes in microtubule dynamics, in particular the onset of catastrophe. We find that our model can reproduce both the experimental microtubule stalling duration and the accompanying EB comet decay rate (Fig 5). At the same time, our model captures the reported mild reduction in catastrophe rate with increasing growth velocity as well as the distribution of stalling durations (Fig. 7AB). Previous 1D models were not able to describe both the mild catastrophe dependence on microtubule growth rates and the size of the stabilizing cap (Brun et al., 2009; Flyvbjerg et al., 1996). A reason for this was the assumption that tubulin dissociation is independent from the microtubule growth velocity (Bowne-Anderson et al., 2013). This necessitated introducing lateral tubulin-tubulin interactions in a 2D model to accurately capture microtubule lifetimes and cap dynamics (Brun et al., 2009; Gardner et al., 2011a). Here, we showed that introducing a highly dynamic tip in a 1D model is sufficient to accurately capture both the microtubule lifetimes as well as the size of the stabilizing cap. The magnitude of the required simulated tip noise would not be observable using fluorescence microscopy, but only with optical tweezers (Gardner et al., 2011a; Kerssemakers et al., 2006; Schek et al., 2007). The extend of any “blurring” of the microtubule tip due to tip noise during frame acquisition would remain below the observable optical resolution (Fig. S6D).

We furthermore showed that our model can capture tubulin washout and reproduces a similar catastrophe delay, tip shrinkage, and comet decay as previously reported (Fig. 6) (Duellberg et al., 2016a). Our model also produces the same correlation between washout delays and growth velocity as was recently observed experimentally (Duellberg et al., 2016a) and thus captures the reported momentary nature of microtubule stability. We conclude that our model can describe tubulin washout and simulate values in good agreement with experiments.

### Microtubule stability depends on the distribution of hydrolysed dimers at the tip

The decay of the EB3 comet during barrier contact can provide further insights into the criterium for microtubule stability. We find that microtubules can remain in a stalled state without the presence of an observable EB3 comet both in our experiments and simulations (compare Fig. 2C and 5D). This suggests that a stalled microtubule does not necessarily require a number of GTP/GDP-Pi subunits that is large enough to be observed as a comet. The effect is explained by the presence of growth fluctuations during microtubule stalling. Hydrolysed subunits at the tip are continually replaced by newly incorporated unhydrolysed subunits, reducing the probability of reaching the critical threshold of *N_unstable_* at the microtubule tip. This phenomenon could also account for reported pausing events during which a microtubule temporarily stops growing without triggering a catastrophe (VanBuren et al., 2005). It illustrates that the onset of a catastrophe is not fully coupled to the presence of an observable comet.

To determine the stretch of hydrolysed subunits at the microtubule tip required to initiate a catastrophe (*N_unstable_*), we measured the ratio between the EB3 comet intensity at the moment of catastrophe and the mean EB3 comet intensity during steady-state growth (Fig. 4BC). In parallel, we evaluated the decay rates of EB3 comets after initial barrier contact. Both measures converge on a catastrophe threshold *N_unstable_* of ~15 uninterrupted hydrolysed terminal subunits, which would approximate a single tubulin layer at the tip of a real 3D microtubule, in line with experimental observations (Caplow and Shanks, 1996; Drechsel and Kirschner, 1994). A recent finding from washout experiments showing that a microtubule requires a stable cap of ~10 tubulin layers at most (Duellberg et al., 2016a) is not at odds with our finding that a catastrophe is triggered when the terminal layer of tubulin is hydrolysed. Because the former result is based on the average remaining density of Mal3 at the moment of catastrophe after tubulin washout, it does not inform on a specific catastrophe criterium. The notion that the stabilizing cap (*L_cap_*) and the EB3 comet (GTP/GDP-Pi region) are large on average, but that only a short stretch of hydrolysed subunits at the microtubule tip is required to trigger a catastrophe, reconciles short and long cap observations (Brun et al., 2009; Duellberg et al., 2016a; Molodtsov et al., 2005; Seetapun et al., 2012; Walker et al., 1991). We thus find that the stability of a microtubule does not primarily depend on the size of the observed EB comet, but instead on the underlying distribution of hydrolysed subunits at the microtubule tip.

### Both the cap size and tip fluctuations determine the onset of catastrophe

To better understand the size of the stabilizing cap (*L_cap_*) and its effect on the catastrophe frequency, we derived an analytical expression for the position of the sequence of hydrolysed subunits equal or greater than *N_unstable_* (see Methods for details). This position is determined by *N_unstable_* and the underlying distribution of hydrolysed subunits. We assume that the density of unhydrolysed dimers decreases exponentially along the microtubule lattice and is fully characterized by *k_hyd_* and 〈*V*〉 (Duellberg et al., 2016a; Maurer et al., 2014; Seetapun et al., 2012). By treating the microtubule lattice as a series of independent Bernoulli trials, we can obtain an expression for the mean position of the first occurrence of a series of hydrolysed subunits equal or greater than *N_unstable_* (Fig. 8A and S5AB). The distance between the microtubule tip and this position is equal to the size of the stabilizing cap, which means we can obtain the relation between the size of the stabilizing cap and the parameters *N_unstable_*, *k_hyd_*, and 〈*V*〉 (Fig. 8BC and S5B). We find that the size of the stabilizing cap scales linearly with the growth velocity 〈*V*〉 (Fig. 8B). The addition of EB3 however affects both the growth velocity and the hydrolysis rate, the combined effect of which results in a decreasing cap size with increasing EB concentration (Fig. 8C). We can calculate the mean size of the stabilizing cap in our model based on *k_hyd_*, 〈*V*〉, and *N_unstable_* and compare it to the length of the GTP/GDP-Pi region for which the EB comet signal is a proxy. Taking 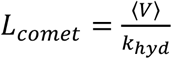 as the characteristic length of the EB comet (Duellberg et al., 2016a; Rickman et al., 2017), we find that the comet underestimates the size of the calculated stabilizing cap (Fig 8C).

**Figure 8.**
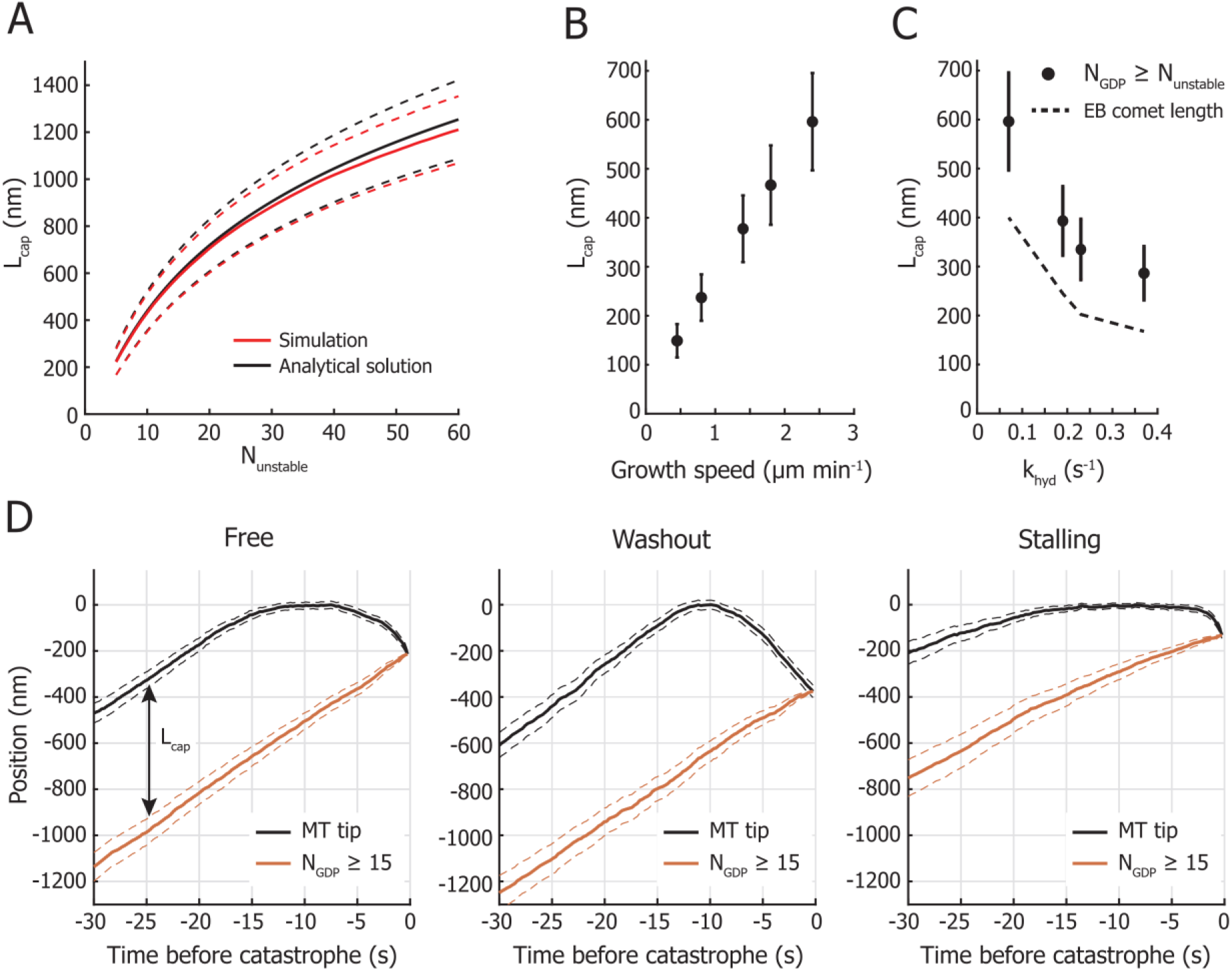
Fluctuations of the stabilizing cap trigger catastrophes. **(A)** Dependence of the cap length *L_cap_* on *N_unstabłe_* (mean ± std). The simulated cap length and the numerical solution based on the analytical model are in good agreement. The shown cap length was determined for the parameters of 0 nM EB3 in Fig. 5C. **(B)** The length of the stabilizing cap *L_cap_* depends linearly on the growth velocity 〈*V*〉. The cap length was calculated with the analytical model (mean ± std) and required a constant *k_hyd_* of 0.1 s^-1^ and an *N_unstable_* of 15, equal to the simulation parameters based on the data from (Janson et al., 2003) (see Fig 7B). **(C)** The mean length of the stabilizing cap *L_cap_* decreases with increasing hydrolysis rate *k_hyd_*. The dependence of the cap size on the hydrolysis rate is calculated with the analytical model (mean ± std) and is based on the parameters of the EB concentrations 0, 20, 50, and 100 nM in Fig 5C. The size of the stabilizing cap based on the position of the sequence *N_unstable_* is larger than the stabilizing cap based on the characteristic length of the EB signal (GTP/GDP-Pi region). **(D)** Simulation of the position of the microtubule tip and of the position of the sequence of hydrolysed subunits defined by *N_unstable_* = 15 prior to catastrophe. Mean traces are for catastrophes during microtubule free growth, after tubulin washout, and during microtubule stalling (mean ± SE). Before the onset of catastrophe, the mean cap length *L_cap_* is constant during steady-state growth as both the position of the tip and the end of the cap move with equal velocity 〈*V*〉. A catastrophe is triggered when *L_cap_* = 0, which is predominantly determined by persistent negative growth excursions of the microtubule tip. All simulations were performed with the parameters for 0 nM EB (Fig. 5C).

Having an expression for the size of the stabilizing cap, we can now explore an intuitive view on how a catastrophe is triggered. The length of the cap is determined by two competing processes on either end, namely noisy growth at the microtubule tip and hydrolysis in the lattice. During steady-state growth, the mean length of the stabilizing cap has a constant size determined by the growth velocity 〈*V*〉 and hydrolysis rate *k_hyd_* (Duellberg et al., 2016a). We observe in our simulations that catastrophes for freely growing microtubules occur as a result of a short period of slowed down or negative growth (Fig 8D left and S3C), in agreement with previous experimental observations (compare Fig. S3 with Fig. 7 in (Maurer et al., 2014)). This shows that the onset of catastrophe is determined by the probability that growth fluctuations remove the stabilizing cap. When we treat the probability for a catastrophe as the probability for negative growth excursions to exceed the length of the stabilizing cap, the microtubule lifetimes are successfully reproduced with the analytical solution (Fig 7A and S5D). This holds true for catastrophes during free growth and after tubulin washout (Fig 8D left and middle). During microtubule stalling however, continuing tip fluctuations replace hydrolysed subunits at the tip for unhydrolysed subunits, reducing the effective mean hydrolysis rate. This results in slowing down of the cap end and delaying the onset of catastrophe (Fig. 8D right). Additionally, the longer catastrophe delay observed with microtubule stalling compared to tubulin washout can be explained by the different effective tip fluctuations. After tubulin washout, loss of the stabilizing cap is caused by both continued hydrolysis and the irreversible loss of tubulin subunits at the tip, whereas the tip of a stalling microtubule can still recover after loss of terminal subunits through tubulin addition and continue to fluctuate (Fig. 8D).

### Microtubule ageing is not required to describe microtubule lifetimes

An observed feature of microtubule stability lacking in our model is an age-dependent catastrophe frequency. It has been reported that “younger” microtubules are more stable than “older” ones (Gardner et al., 2011b; Odde et al., 1995). Ageing has also been observed through a gradual reduction of the EB comet intensity during steady-state growth (Maurer et al., 2012; Mohan et al., 2013) and through shorter catastrophe delays after tubulin washout for older microtubules (Duellberg et al., 2016b). The two proposed processes responsible for inferring ageing are either based on a multi-step lattice defect model (Bowne-Anderson et al., 2013; Mohan et al., 2013) or on tapering of the microtubule tip during growth (Chretien et al., 1995; Coombes et al., 2013; Duellberg et al., 2016b; VanBuren et al., 2005). Invariably, these mechanisms are tightly coupled to the onset of a catastrophe and are required to understand microtubule lifetimes. In our model however, we find that the onset of a catastrophe is independent from microtubule ageing.

Microtubule ageing is generally characterized by fitting lifetime distributions to a Gamma distribution to obtain the shape parameter, a measure for the number of sequential steps required to trigger a catastrophe (Gardner et al., 2011b; Odde et al., 1995). In our experimental lifetime distribution, we find that the Gamma shape parameter is independent from the EB concentration (Fig. S6A), in line with previous reports (Mohan et al., 2013). To account for microtubule ageing in our model, we introduced time-dependent tip fluctuations (Fig. S6B). Increasing the growth fluctuations with time while keeping the mean growth velocity constant, increases the probability of reducing the stabilizing cap to zero due to an increase in negative growth excursions. By increasing *D_tip_* with a rate of 0.025 s^-1^ following a bounded exponential curve, we can reproduce the experimental ageing parameters (Fig. S6C). Linking microtubule ageing to an increase in tip noise and not to the hydrolysis rate or to the accumulation of defects in the microtubule lattice also agrees with previous studies (Zakharov et al., 2015). It was reported that microtubule ageing is in fact correlated with the frequency of encountering curled protofilaments (Zakharov et al., 2015), an effect that can be represented with our phenomenological description of increasing tip noise (McIntosh et al., 2018). So, although microtubule ageing is not required to accurately capture microtubule lifetimes, it can be easily incorporated by introducing time-dependent tip fluctuations.

### Outlook

Our experimental setup will be useful for studying microtubule interactions and the functional effect of stabilizing and destabilizing microtubule associated proteins. Our approach can be used to study the influence of MAPs (Meadows et al., 2018), tubulin isotypes and PTMs (Sirajuddin et al., 2014) on the stability of pushing microtubules. Furthermore, the SiC overhangs are designed to be compatible with a previously published method to specifically functionalize the barriers with protein complexes (Taberner et al., 2014), enabling the study of microtubule end-on interactions with TIRF microscopy (Vleugel et al., 2016).

A possible extension of our 1D model to describe the effect of +TIPs on microtubule dynamics in general could be to characterize them phenomenologically by their effect on GTP hydrolysis and tip fluctuations. As the effect of EB3 can be described this way, we hypothesize that the effect of other microtubule associated proteins can be characterized similarly. To allow future extensions and modifications of our model, we have made the simulation source code available under an open-source licence on GitHub (https://github.com/florian-huber/mtdynamics) with extra documentation.

## METHODS AND MATERIALS

### Proteins

GFP-EB3 was a kind gift from Michel Steinmetz. All tubulin products were acquired from Cytoskeleton Inc, with all unlabelled tubulin specifically from a single lot.

### Microfabrication of barriers

The fabrication method for the micro-fabricated barriers with an SiC overhang is inspired by (Kalisch et al., 2011), (Taberner et al., 2014), and (Aher et al., 2018). All fabrication steps were performed in a cleanroom environment (van Leeuwenhoek Laboratory, NanoLab NL). The barrier was designed with the following considerations in mind:

- The width of the channels should favour stalling events over buckling events, but remain large enough for GMPCPP-stabilized seeds to easily land.
- A bottom layer of SiC is needed to prevent etching into the coverslip during a Buffered Oxide Etch. This layer needs to be as thin as possible to prevent photon absorption by the semiconductor resulting in a diminished signal-to-noise and surface heating. Although SiC is transparent for wavelengths > 0.5 μm, its bandgap of ~2.8 eV can result in photon absorption for the commonly used 405 and 488 nm lasers (Pham, 2004). Using PE-CVD, 10 nm is thinnest layer we could fabricate while still maintaining the layer’s integrity to protect the coverslip from the Buffered Oxide Etch.
- The layer of SiO_2_ of 100 nm ensures that the microtubule can polymerize underneath the overhang while remaining inside the evanescent wave.
- The top layer of SiC is 250 nm thick to ensure mechanical stability, while still allowing to observe microtubules growing on top of the barrier despite some photon absorption.

To start, glass coverslips (24×24 mm, #1) were cleaned for 10 min with base piranha, a mixture of H_2_O:NH_4_OH:H_2_O_2_ in a 5:1:1 ratio heated to 70°C. Then, three sequential layers of SiC (10 nm), SiO_2_ (100 nm), and SiC (250 nm) are deposited on the cleaned surface via Plasma-Enhanced Chemical Vapour Deposition (PE-CVD) at 300°C (Oxford Instruments PlasmaPro 80). PE-CVD ensures a surface smooth enough for TIRF microscopy with fast deposition rates (70 nm/min for SiO_2_ and 40 nm/min for SiC).

In order to transfer the barrier pattern to the surface, UV lithography is used. First, to aid in the adhesion of the photoresist, a few drops of hexamethyldisilazane (HMDS) are spin coated on the SiC surface and allowed to dry on a 115°C hotplate for 30 seconds. Then a 1.3 μm layer of the positive photoresist S1813 (MicroChem) is spin coated (5000rpm) on the surface and pre-baked for 90 seconds on a 115°C hotplate. Exposure of the photoresist through a chromium mask with a near-UV source (320-365 nm, approx. 13 mW/cm^2^) transfers the barrier pattern in 4 seconds (EVgroup EVG 620). Development with MF321 (MicroPosit) for 60 seconds removes the UV-exposed regions of the resist.

Next, Reactive Ion Etching (Leybold Hereaus) with a mixture of CHF_3_:O_2_ (50 sccm:2.5 sccm) etches through the exposed regions of the 250 nm SiC layer and into the SiO_2_ layer. The etch is performed at 50 μbar and at 100 W, resulting in a bias voltage of 400 V. It is important to etch completely through the top SiC layer, but only partly through the SiO_2_ layer, to leave the bottom SiC layer intact. Any remaining photoresist after the etch is removed by sonication of the sample in acetone for 10 minutes.

Finally, the sample is submerged in buffered hydrofluoric acid (HF:NH_4_F = 12.5:87.5%) to selectively etch the exposed SiO_2_ with a rate of approximately 200 nm/min to obtain an overhang of 1.5 μm. The final barriers are 100 nm high with an overhang of 1.5 μm, enclosing channels with a width of 15 μm.

### *In vitro* microtubule dynamics assay

Reconstitution of microtubule dynamics was performed as previously described in (Bieling et al., 2007; Montenegro Gouveia et al., 2010). After cleaning the barrier sample with O_2_-plasma, a flow channel was constructed with a cleaned glass slide and double-sided sticky tape in such a way that the channel direction is perpendicular to the barriers. Then, the surface was consecutively functionalized with 0.5 mg/ml PLL-PEG-biotin(20%) (SuSoS AG, Switzerland), 0.2 mg/ml NeutrAvidin (Invitrogen), and 0.5 mg/ml κ-casein (Sigma). All components were kept in MRB80 buffer, comprised of 80mM piperazine-N,N’-bis(2-ethanesulfonic acid), 4mM MgCl_2_, and 1mM EGTA at a pH of 6.8. The reaction mixture contained 15 μM tubulin (7% rhodamine labelled) in the presence of GFP-EB3 or Hilyte488 labelled tubulin in the absence of GFP-EB3, supplemented with 0.5 mg/ml κ-casein, 0.15% methylcellulose, 50 mM KCl, 1 mM GTP, oxygen scavenger mix (4 mM DTT, 200 μg/ml catalase, 400 μg/ml glucose oxidase, 50mM glucose). The reaction mix is then centrifuged in an Airfuge (Beckman Coulter) at 30psi for 8 minutes to remove any aggregated complexes before being introduced to the sample. GMPCPP-stabilized seeds (70% unlabelled tubulin, 18% biotinylated tubulin, 12% rhodamine-labelled tubulin) were introduced to the channel with the flow direction perpendicular to the barriers. Flow cells were sealed with vacuum grease and imaged on a TIRF microscope at 28-30°C.

### TIRF microscopy

All experiments were imaged using TIRF microscopy, consisting of an Ilas^2^ system (Roper Scientific) on a Nikon Ti-E inverted microscope. The Ilas^2^ system is a dual illuminator for azimuthal spinning TIRF illumination equipped with a 150 mW 488 nm laser, a 100 mW 561 nm laser, and a ZT405/488/561/640rpc dichroic mirror. Simultaneous dual-acquisition was performed with two Evolve 512 EMCCD camera’s (Photometrics) through a 525/50 nm and a 609/54 emission filter, using a Nikon CFI Plan Apochromat 100XH NA1.45 TIRF oil objective. Together with an additional magnifying lens, the final magnification resulted in a pixel size of 107 nm/pixel. The sample was heated with a custom objective heater to 28-30°C and was kept in focus with the Nikon Perfect Focus system. The hardware was controlled with MetaMorph 7.8.8.0 (Molecular Device).

### Image treatment

The image stacks obtained with TIRF microscopy were corrected prior to data analysis. First, simultaneous acquisition of rhodamine-labelled tubulin and GFP-EB3 on two cameras introduced a non-linear spatial offset between the two image stacks due to imperfections in the dichroic mirror and in the alignment of the two cameras. By scanning multiple FOVs of a calibration slide containing 100 nm TetraSpeck beads (ThermoFisher) and automatically locating the centroids through a custom written MATLAB script, a non-linear registration profile accounting for the spatial offset was calculated. The misaligned image stack was corrected by applying this registration profile based on the position of ~500 bead positions. Additionally, any sample drift was corrected by subpixel image registration through cross-correlation (Guizar-Sicairos et al., 2008).

Secondly, some scattering of excitation light at the edge of the SiC overhang made proper determination of the GFP-EB3 signal near the barrier difficult. Although this effect was mostly mediated by creating a wide undercut that physically separated the edge of the overhang from the barrier, a correction was nonetheless applied. To correct the signal, the minimum intensity value of each pixel in the image stack was subtracted from that pixel in each image. This correction enabled tracking of the EB3 comet near the barrier and accurate measurement of the EB3 comet intensity.

Thirdly, a general background subtraction was performed in Fiji (Schindelin et al., 2012) to correct for inhomogeneous illumination.

### Image analysis

Analysis of the images was partly performed with Fiji and with MATLAB. After the image treatment described above, kymographs were created by drawing straight lines of 9-pixel width (0.95 μm) along growing microtubules using KymoResliceWide plugin with maximum transverse intensity (http://fiji.sc/KymoResliceWide). Each growth event in the kymographs was manually traced to determine the position of the microtubule tip. This position was then used to fit the EB3 comet to obtain its position and intensity, using the intensity profile:

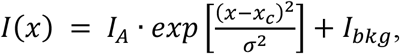

where *I*(*x*) is the fluorescence intensity, *I_bkg_* is the background intensity, *I_A_* is the intensity amplitude, *x_c_* is the position of the peak of the EB3 comet, and *σ* is the width of the EB3 comet. As EB3 comet decay at the barrier makes fitting impossible, the intensity during contact was determined by calculating the average intensity value in a region around the comet position and around the barrier (Fig. S1C). The barrier contact duration and comet decay duration were determined manually.

### Monte Carlo simulations

Simulations of growing microtubules were run as a series of discrete, fixed time-steps. The length of the time steps *δt* was chosen small enough to properly account for the random hydrolysis of the subunits (*P_hyd_*, the probability for a dimer to undergo hydrolysis within one timestep was kept at <0.05). Further restrictions were to not exceed the desired framerate, in our case the lowest used experimental frame rate of 250 ms. Due to the discrete nature of microtubule growth in subunits, the next-lowest time-step for which 〈*V*〉*δt*/*L*_0_ became an integer was chosen, with 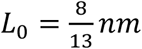 the length increment per subunit and 〈*V*〉 the microtubule mean growth velocity.

Each microtubule simulation started from a few initial subunits (a ‘seed’) that were excluded from hydrolysis, and that were not allowed to be removed during microtubule tip fluctuations. Microtubule growth was simulated as a discrete, biased, Gaussian random walk. This means that for each time-step *δt*, the microtubule length was changed by a discretized random number of subunits that was drawn from a Gaussian distribution with standard deviation 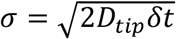 and centered at 〈*dx*〉/*L*_0_.

During each time step, subunits transition from the GTP/GDP-Pi to the GDP state by random hydrolysis with a rate *k_hyd_*. Whenever the foremost uninterrupted strand of GDP state subunits (≥ *N_unstable_* subunits in a row) is changed, the position of the end of the stable cap will jump to the front element of this strand, which we interpret as the new position of the end of the stable cap *L_end–of–cap_*.

A simulation run ends when a catastrophe occurs. This happens when the stable cap shrinks to zero, i.e. if *L_tip_* − *L_end–of–cap_* = 0, where *L_tip_* is defined as the position of the foremost subunit of the microtubule. The growth duration was defined as the time from initial growth until catastrophe. To exclude nucleation kinetics from the simulated lifetimes, a microtubule is considered to grow after reaching a length of 250 nm.

The presence of a physical barrier is modelled by introducing a fixed barrier position *L_barrier_*. Tip dynamics and random hydrolysis remained unchanged, only the microtubule length was truncated whenever it would penetrate the barrier. This means the length of the microtubule was set back to *L_barrier_* if *L_tip_* > *L_barrier_*. The barrier contact time was then defined as the time from the microtubules first contact with the barrier until its catastrophe.

The simulation was written in Python 3.6 and run on standard PCs. The code to run the simulation is available under an open-license on GitHub (https://github.com/florian-huber/mtdynamics).

### Analytical expression for the size of the stabilizing cap

The length of the stabilizing cap *L_cap_* is defined as the distance between the position of the microtubule tip and the first occurrence of a sequence of hydrolysed subunits *N* equal or greater than *N_unstable_* (Fig. 3C). The location of this sequence of hydrolysed subunits is determined by the distribution of GDP dimers in the microtubule lattice. We assume that the GTP/GDP-Pi distribution at the microtubule tip decays mono-exponentially and depends on the hydrolysis rate *k_hyd_* and mean growth velocity 〈*V*〉 (Bieling et al., 2007; Duellberg et al., 2016a; Seetapun et al., 2012). The probability *p*(*x*) of finding a GTP/GDP-Pi subunit at position *x* in the lattice (with the microtubule tip at *x* = 0) corresponds to

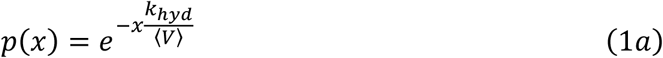

with the probability of finding a GDP subunit at position *x* being

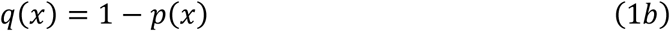

To find the probability distribution of the position of a sequence of *N* sequential hydrolysed dimers equal or greater than *N_unstable_*, we treat the discrete 1D lattice as a series of independent Bernoulli trials with probabilities *p*(*x*) and *q*(*x*) for GTP/GDP-Pi or GDP dimers respectively (Fig. S5A). For a lattice shorter than *N*, the probability of finding a sequence of *N* GDP is zero since the sequence is longer than the considered lattice. The probability of finding *N* GDP subunits between the positions *x* = 1 and *x* = *N* is equal to the product of the probability at each position *x*:

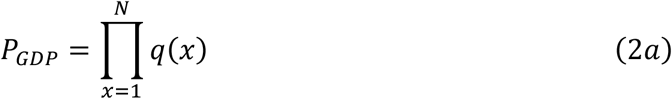

If *N* and 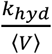 and are small, the probability of finding a GDP subunit at the beginning of the sequence is approximately equal to that at the end, i.e. *q*(*x*_1_) ≈ *q*(*x_N_*). Using this assumption, equation (2*a*) can be rewritten as

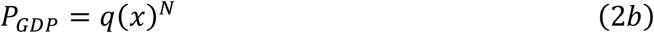

For a position on the lattice further from the tip, the probability of finding a sequence *N* ≥ *N_unstable_* at position *x*, with *x* being the first position of the sequence, is equal to

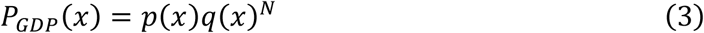

as the dimer directly preceding the sequence needs to be unhydrolyzed to initiate the sequence (Fig S5A).

We can now obtain an expression for the probability of finding this sequence for <u>the first time</u> at position *x*, by considering the probability that no sequence is found at any position closer to the microtubule tip:

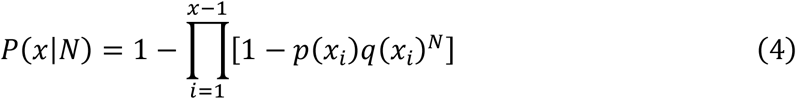

This expression gives the cumulative distribution for finding a sequence of *N* GDP subunits at position *x* during steady-state growth. We find that this approximation holds reasonably well for the entire range of *N_unstable_* we explored using the 1D simulation (Fig. S5B). The probability of finding this sequence of GDP can be captured by the Gaussian cumulative distribution function:

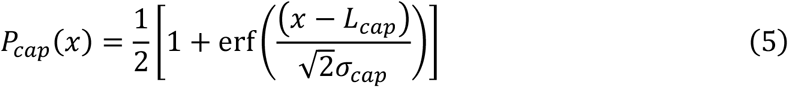

Through numerical analysis we find that the dependency of *L_cap_* on *N* follows a power law (Fig. 8A) and can be described with

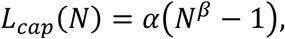

where *α* and *β* are coefficients that depend on the hydrolysis rate and the growth velocity. Similarly, we can calculate the dependence of the cap size on the parameters *k_hyd_* and 〈*V*〉 (Fig. 8BC).

### Analytical expression for the catastrophe probability and microtubule lifetimes

The probability for a microtubule to undergo a catastrophe within time window Δ*t* is defined as *P_cat_*(Δ*t*) and is equal to the probability of reducing the cap size *L_cap_* to zero during Δ*t*. The cap size evolves by two competing stochastic processes: it increases by dimer addition at the tip and shrinks by dimer removal from the tip and hydrolysis in the lattice. To obtain an analytical expression for the catastrophe probability, we consider a microtubule at steady-state growth. In this frame of reference, the end of the cap is on average a constant distance from the microtubule tip, as both the microtubule tip and the position of the cap end move with equal velocity 〈*V*〉. Any fluctuations of the cap size during steady-state growth are caused by fluctuations of the tip position. However, due to the stochastic nature of hydrolysis, any incorporated dimers at the microtubule tip position only affect the position of the cap end after a characteristic time delay *τ_c_*, which is approximately equal to *k_hyd_*^−1^ (Fig. S5C). In other words, the delay gives a measure of the time window during which the fluctuations can affect *L_cap_*, before the position of the cap is affected by hydrolysis. The catastrophe probability is then equal to the probability of tip fluctuations to exceed the position of *L_cap_* during time window *τ_c_*.

The growth fluctuations at the microtubule tip can be described by a biased random walk with Gaussian distributed steps Δ*x* within Δ*t* (Fig 3B).

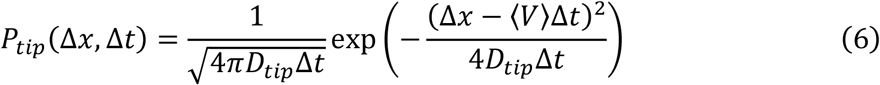

Note that in the frame of reference of steady-state growth, 〈*V*〉 = 0. To find *P_cat_*(Δ*t*), we calculate the probability that the microtubule tip exceeds *L_cap_* during Δ*t* ≤ *τ_c_*. The survival probability, the probability that the microtubule tip does not exceed *L_cap_* for all times up to *τ_c_*, is defined as

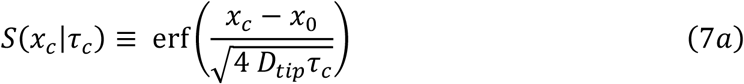

with *x_c_* being the critical cap-end position. Since the critical cap-end position is defined with respect to *x*_0_ being the moving tip, we can set *x*_0_ = 0 and get the probability for the tip to have reached *x* ≥ *x_c_*:

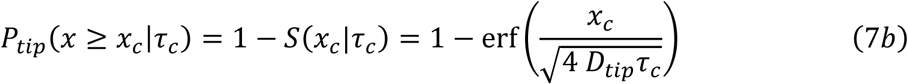

When we set *x_c_* = *L_cap_*, we can calculate the catastrophe probability *P_cat_*(*τ_c_*) as the probability of the tip fluctuations exceeding the cap size *L_cap_* during time window *τ_c_* with

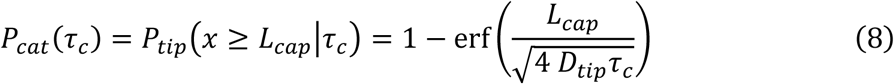

The microtubule lifetime distribution *T_cat_*(*t*) can then be obtained by calculating the fraction of microtubules that underwent a catastrophe after each timestep *τ_c_* (Fig. 7A and S5D):

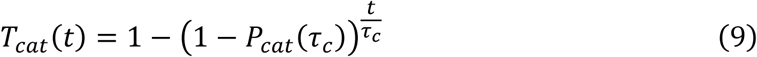

Note that the steady-state approximation becomes less accurate for large 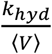 as steady-state growth might not be reached in the first place. This leads to a lack of short events and consequently to an overestimation of the microtubule lifetimes.

## Author contributions

M.K. designed and fabricated the micro-fabricated barriers, performed experiments with EB and analysed the data, developed the analytical solution, and ran simulations. F.H. developed the 1D microtubule model, wrote the simulation code and ran and analysed the simulations. SM.K. performed and analysed the experiments with Mal3. M.K, F.H., and M.D. wrote the paper. M.D. coordinated the project.

## Acknowledgements

We thank M.O. Steinmetz for the gift of GFP-EB3 and the technicians of the van Leeuwenhoek Laboratory for their technical assistance in the fabrication of the barriers. We thank Vladimir Volkov and Louis Reese for discussions and critical reading of the manuscript and Seungkyu Ha for his assistance in acquiring the SEM images. This work was supported by the European Research Council Synergy grant 6098822 to Marileen Dogterom.

## REFERENCES

Aher, A., M. Kok, A. Sharma, A. Rai, N. Olieric, R. Rodriguez-Garcia, E.A. Katrukha, T. Weinert, V. Olieric, L.C. Kapitein, M.O. Steinmetz, M. Dogterom, and A. Akhmanova. 2018. {CLASP} Suppresses Microtubule Catastrophes through a Single {TOG} Domain. Dev Cell. 46:40–58.e48, https://dx.doi.org/10.1016/j.devcel.2018.05.032

Akhmanova, A., and M.O. Steinmetz. 2015. Control of microtubule organization and dynamics: two ends in the limelight. Nat Rev Mol Cell Biol. 16:711–726, https://dx.doi.org/10.1038/nrm4084

Alushin, G.M., G.C. Lander, E.H. Kellogg, R. Zhang, D. Baker, and E. Nogales. 2014. High-resolution microtubule structures reveal the structural transitions in αβ-tubulin upon {GTP} hydrolysis. Cell. 157:1117–1129, https://dx.doi.org/10.1016/j.cell.2014.03.053

Antal, T., P.L. Krapivsky, S. Redner, M. Mailman, and B. Chakraborty. 2007. Dynamics of an idealized model of microtubule growth and catastrophe. Phys Rev E Stat Nonlin Soft Matter Phys. 76:041907, https://dx.doi.org/10.1103/PhysRevE.76.041907

Bayley, P., M. Schilstra, and S. Martin. 1989. A lateral cap model of microtubule dynamic instability. FEBS Lett. 259:181–184, https://dx.doi.org/10.1016/0014-5793(89)81523-6

Bieling, P., L. Laan, H. Schek, E.L. Munteanu, L. Sandblad, M. Dogterom, D. Brunner, and T. Surrey. 2007. Reconstitution of a microtubule plus-end tracking system in vitro. Nature. 450:1100–1105, https://dx.doi.org/10.1038/nature06386

Bollinger, J.A., and M.J. Stevens. 2019. Diverse balances of tubulin interactions and shape change drive and interrupt microtubule depolymerization. Soft Matter. 15:8137–8146, https://dx.doi.org/10.1039/c9sm01323g

Bouchet, B.P., I. Noordstra, M. van Amersfoort, E.A. Katrukha, Y.C. Ammon, N.D. Ter Hoeve, L. Hodgson, M. Dogterom, P.W. Derksen, and A. Akhmanova. 2016. Mesenchymal Cell Invasion Requires Cooperative Regulation of Persistent Microtubule Growth by {SLAIN}2 and {CLASP}1. Dev Cell. 39:708–723, https://dx.doi.org/10.1016/j.devcel.2016.11.009

Bowne-Anderson, H., M. Zanic, M. Kauer, and J. Howard. 2013. Microtubule dynamic instability: a new model with coupled {GTP} hydrolysis and multistep catastrophe. Bioessays. 35:452–461, https://dx.doi.org/10.1002/bies.201200131

Brangwynne, C.P., F.C. MacKintosh, S. Kumar, N.A. Geisse, J. Talbot, L. Mahadevan, K.K. Parker, D.E. Ingber, and D.A. Weitz. 2006. Microtubules can bear enhanced compressive loads in living cells because of lateral reinforcement. J Cell Biol. 173:733–741, https://dx.doi.org/10.1083/jcb.200601060

Brangwynne, C.P., F.C. MacKintosh, and D.A. Weitz. 2007. Force fluctuations and polymerization dynamics of intracellular microtubules. Proc Natl Acad Sci U S A. 104:16128–16133, https://dx.doi.org/10.1073/pnas.0703094104

Brouhard, G.J., and L.M. Rice. 2018. Microtubule dynamics: an interplay of biochemistry and mechanics. Nat Rev Mol Cell Biol. 19:451–463, https://dx.doi.org/10.1038/s41580-018-0009-y

Brun, L., B. Rupp, J.J. Ward, and F. Nedelec. 2009. A theory of microtubule catastrophes and their regulation. Proc Natl Acad Sci U S A. 106:21173–21178, https://dx.doi.org/10.1073/pnas.0910774106

Caplow, M., and J. Shanks. 1996. Evidence that a single monolayer tubulin-{GTP} cap is both necessary and sufficient to stabilize microtubules. Mol Biol Cell. 7:663–675, https://dx.doi.org/10.1091/mbc.7.4.663

Carlier, M.F., and D. Pantaloni. 1981. Kinetic analysis of guanosine 5′-triphosphate hydrolysis associated with tubulin polymerization. Biochemistry. 20:1918–1924, http://www.ncbi.nlm.nih.gov/pubmed/7225365

Carlier, M.F., and D. Pantaloni. 1982. Assembly of microtubule protein: role of guanosine di- and triphosphate nucleotides. Biochemistry. 21:1215–1224, https://www.ncbi.nlm.nih.gov/pubmed/7074077

Chen, Y.D., and T.L. Hill. 1985. Monte {C}arlo study of the {GTP} cap in a five-start helix model of a microtubule. Proc Natl Acad Sci U S A. 82:1131–1135, https://www.ncbi.nlm.nih.gov/pubmed/3856250

Chretien, D., S.D. Fuller, and E. Karsenti. 1995. Structure of growing microtubule ends: two-dimensional sheets close into tubes at variable rates. J Cell Biol. 129:1311–1328, https://dx.doi.org/10.1083/jcb.129.5.1311

Coletti, C., M.J. Jaroszeski, A. Pallaoro, A.M. Hoff, S. Iannotta, and S.E. Saddow. 2007. Biocompatibility and wettability of crystalline {S}i{C} and {S}i surfaces. Conf Proc IEEE Eng Med Biol Soc. 2007:5850–5853, https://dx.doi.org/10.1109/IEMBS.2007.4353678

Colin, A., P. Singaravelu, M. Thery, L. Blanchoin, and Z. Gueroui. 2018. Actin-Network Architecture Regulates Microtubule Dynamics. Curr Biol, https://dx.doi.org/10.1016/j.cub.2018.06.028

Coombes, C.E., A. Yamamoto, M.R. Kenzie, D.J. Odde, and M.K. Gardner. 2013. Evolving tip structures can explain age-dependent microtubule catastrophe. Curr Biol. 23:1342–1348, https://dx.doi.org/10.1016/j.cub.2013.05.059

Das, D., D. Das, and R. Padinhateeri. 2014. Force-induced dynamical properties of multiple cytoskeletal filaments are distinct from that of single filaments. PLoS One. 9:e114014, https://dx.doi.org/10.1371/journal.pone.0114014

Debs, G.E., M. Cha, X. Liu, A.R. Huehn, and C.V. Sindelar. 2020. Dynamic and asymmetric fluctuations in the microtubule wall captured by high-resolution cryoelectron microscopy. Proc Natl Acad Sci U S A. 117:16976–16984, https://dx.doi.org/10.1073/pnas.2001546117

Dhar, S., O. Seitz, M.D. Halls, S. Choi, Y.J. Chabal, and L.C. Feldman. 2009. Chemical properties of oxidized silicon carbide surfaces upon etching in hydrofluoric acid. J Am Chem Soc. 131:16808–16813, https://dx.doi.org/10.1021/ja9053465

Dogterom, M., and G.H. Koenderink. 2019. Actin-microtubule crosstalk in cell biology. Nat Rev Mol Cell Biol. 20:38–54, https://dx.doi.org/10.1038/s41580-018-0067-1

Drechsel, D.N., and M.W. Kirschner. 1994. The minimum GTP cap required to stabilize microtubules. Curr Biol. 4:1053–1061, https://dx.doi.org/10.1016/s0960-9822(00)00243-8

Duellberg, C., N.I. Cade, D. Holmes, and T. Surrey. 2016a. The size of the {EB} cap determines instantaneous microtubule stability. Elife. 5, https://dx.doi.org/10.7554/eLife.13470

Duellberg, C., N.I. Cade, and T. Surrey. 2016b. Microtubule aging probed by microfluidics-assisted tubulin washout. Mol Biol Cell. 27:3563–3573, https://dx.doi.org/10.1091/mbc.E16-07-0548

Flyvbjerg, H., T.E. Holy, and S. Leibler. 1996. Microtubule dynamics: {C}aps, catastrophes, and coupled hydrolysis. Phys Rev E Stat Phys Plasmas Fluids Relat Interdiscip Topics. 54:5538–5560, https://www.ncbi.nlm.nih.gov/pubmed/9965740

Gardner, M.K., B.D. Charlebois, I.M. Janosi, J. Howard, A.J. Hunt, and D.J. Odde. 2011a. Rapid microtubule self-assembly kinetics. Cell. 146:582–592, https://dx.doi.org/10.1016/j.cell.2011.06.053

Gardner, M.K., M. Zanic, C. Gell, V. Bormuth, and J. Howard. 2011b. Depolymerizing kinesins {K}ip3 and {MCAK} shape cellular microtubule architecture by differential control of catastrophe. Cell. 147:1092–1103, https://dx.doi.org/10.1016/j.cell.2011.10.037

Gregoretti, I.V., G. Margolin, M.S. Alber, and H.V. Goodson. 2006. Insights into cytoskeletal behavior from computational modeling of dynamic microtubules in a cell-like environment. J Cell Sci. 119:4781–4788, https://dx.doi.org/10.1242/jcs.03240

Guizar-Sicairos, M., S.T. Thurman, and J.R. Fienup. 2008. Efficient subpixel image registration algorithms. Opt Lett. 33:156–158, https://www.ncbi.nlm.nih.gov/pubmed/18197224

Gurel, P.S., A.L. Hatch, and H.N. Higgs. 2014. Connecting the cytoskeleton to the endoplasmic reticulum and {G}olgi. Curr Biol. 24:R660–R672, https://dx.doi.org/10.1016/j.cub.2014.05.033

Janson, M.E., M.E. de Dood, and M. Dogterom. 2003. Dynamic instability of microtubules is regulated by force. J Cell Biol. 161:1029–1034, https://dx.doi.org/10.1083/jcb.200301147

Kalisch, S.M., L. Laan, and M. Dogterom. 2011. Force generation by dynamic microtubules in vitro. Methods Mol Biol. 777:147–165, https://dx.doi.org/10.1007/978-1-61779-252-6_11

Karr, T.L., and D.L. Purich. 1978. Examination of tubulin-nucleotide interactions by protein fluorescence quenching measurements. Biochem Biophys Res Commun. 84:957–961, https://www.ncbi.nlm.nih.gov/pubmed/728162

Kerssemakers, J.W., E.L. Munteanu, L. Laan, T.L. Noetzel, M.E. Janson, and M. Dogterom. 2006. Assembly dynamics of microtubules at molecular resolution. Nature. 442:709–712, https://dx.doi.org/10.1038/nature04928

Kim, T., and L.M. Rice. 2019. Long-range, through-lattice coupling improves predictions of microtubule catastrophe. Mol Biol Cell. 30:1451–1462, https://dx.doi.org/10.1091/mbc.E18-10-0641

Komarova, Y.A., I.A. Vorobjev, and G.G. Borisy. 2002. Life cycle of {MT}s: persistent growth in the cell interior, asymmetric transition frequencies and effects of the cell boundary. J Cell Sci. 115:3527–3539, https://www.ncbi.nlm.nih.gov/pubmed/12154083

Laan, L., J. Husson, E.L. Munteanu, J.W. Kerssemakers, and M. Dogterom. 2008. Force-generation and dynamic instability of microtubule bundles. Proc Natl Acad Sci U S A. 105:8920–8925, https://dx.doi.org/10.1073/pnas.0710311105

Lee, C.T., and E.M. Terentjev. 2019. Structural effects of cap, crack, and intrinsic curvature on the microtubule catastrophe kinetics. J Chem Phys. 151:135101, https://dx.doi.org/10.1063/1.5122304

Letort, G., F. Nedelec, L. Blanchoin, and M. Thery. 2016. Centrosome centering and decentering by microtubule network rearrangement. Mol Biol Cell. 27:2833–2843, https://dx.doi.org/10.1091/mbc.E16-06-0395

Margolin, G., I.V. Gregoretti, T.M. Cickovski, C. Li, W. Shi, M.S. Alber, and H.V. Goodson. 2012. The mechanisms of microtubule catastrophe and rescue: implications from analysis of a dimer-scale computational model. Mol Biol Cell. 23:642–656, https://dx.doi.org/10.1091/mbc.E11-08-0688

Margolin, G., I.V. Gregoretti, H.V. Goodson, and M.S. Alber. 2006. Analysis of a mesoscopic stochastic model of microtubule dynamic instability. Phys Rev E Stat Nonlin Soft Matter Phys. 74:041920, https://dx.doi.org/10.1103/PhysRevE.74.041920

Maurer, S.P., P. Bieling, J. Cope, A. Hoenger, and T. Surrey. 2011. {GTP}γ{S} microtubules mimic the growing microtubule end structure recognized by end-binding proteins ({EB}s). Proc Natl Acad Sci U S A. 108:3988–3993, https://dx.doi.org/10.1073/pnas.1014758108

Maurer, S.P., N.I. Cade, G. Bohner, N. Gustafsson, E. Boutant, and T. Surrey. 2014. {EB}1 accelerates two conformational transitions important for microtubule maturation and dynamics. Curr Biol. 24:372–384, https://dx.doi.org/10.1016/j.cub.2013.12.042

Maurer, S.P., F.J. Fourniol, G. Bohner, C.A. Moores, and T. Surrey. 2012. {EB}s recognize a nucleotide-dependent structural cap at growing microtubule ends. Cell. 149:371–382, https://dx.doi.org/10.1016/j.cell.2012.02.049

McIntosh, J.R., E. O’Toole, G. Morgan, J. Austin, E. Ulyanov, F. Ataullakhanov, and N. Gudimchuk. 2018. Microtubules grow by the addition of bent guanosine triphosphate tubulin to the tips of curved protofilaments. J Cell Biol, https://dx.doi.org/10.1083/jcb.201802138

Meadows, J.C., L.J. Messin, A. Kamnev, T.C. Lancaster, M.K. Balasubramanian, R.A. Cross, and J.B. Millar. 2018. Opposing kinesin complexes queue at plus tips to ensure microtubule catastrophe at cell ends. EMBO Rep. 19, https://dx.doi.org/10.15252/embr.201846196

Michaels, T.C., S. Feng, H. Liang, and L. Mahadevan. 2020. Mechanics and kinetics of dynamic instability. Elife. 9, https://dx.doi.org/10.7554/eLife.54077

Mitchison, T., and M. Kirschner. 1984. Dynamic instability of microtubule growth. Nature. 312:237–242, http://www.ncbi.nlm.nih.gov/pubmed/6504138

Mohan, R., E.A. Katrukha, H. Doodhi, I. Smal, E. Meijering, L.C. Kapitein, M.O. Steinmetz, and A. Akhmanova. 2013. End-binding proteins sensitize microtubules to the action of microtubule-targeting agents. Proc Natl Acad Sci U S A. 110:8900–8905, https://dx.doi.org/10.1073/pnas.1300395110

Molodtsov, M.I., E.A. Ermakova, E.E. Shnol, E.L. Grishchuk, J.R. McIntosh, and F.I. Ataullakhanov. 2005. A molecular-mechanical model of the microtubule. Biophys J. 88:3167–3179, https://dx.doi.org/10.1529/biophysj.104.051789

Montenegro Gouveia, S., K. Leslie, L.C. Kapitein, R.M. Buey, I. Grigoriev, M. Wagenbach, I. Smal, E. Meijering, C.C. Hoogenraad, L. Wordeman, M.O. Steinmetz, and A. Akhmanova. 2010. In vitro reconstitution of the functional interplay between {MCAK} and {EB}3 at microtubule plus ends. Curr Biol. 20:1717–1722, https://dx.doi.org/10.1016/j.cub.2010.08.020

Nguyen-Ngoc, T., K. Afshar, and P. Gonczy. 2007. Coupling of cortical dynein and {G}α proteins mediates spindle positioning in {C}aenorhabditis elegans. Nat Cell Biol. 9:1294–1302, https://dx.doi.org/10.1038/ncb1649

Nogales, E. 1999. A structural view of microtubule dynamics. Cell Mol Life Sci. 56:133–142, http://www.ncbi.nlm.nih.gov/pubmed/11213253

Odde, D.J., L. Cassimeris, and H.M. Buettner. 1995. Kinetics of microtubule catastrophe assessed by probabilistic analysis. Biophys J. 69:796–802, https://dx.doi.org/10.1016/S0006-3495(95)79953-2

Odde, D.J., L. Ma, A.H. Briggs, A. DeMarco, and M.W. Kirschner. 1999. Microtubule bending and breaking in living fibroblast cells. J Cell Sci. 112 ( Pt 19):3283–3288, https://www.ncbi.nlm.nih.gov/pubmed/10504333

Ohi, R., and M. Zanic. 2016. Ahead of the Curve: New Insights into Microtubule Dynamics. F1000Res. 5, https://dx.doi.org/10.12688/f1000research.7439.1

Padinhateeri, R., A.B. Kolomeisky, and D. Lacoste. 2012. Random hydrolysis controls the dynamic instability of microtubules. Biophys J. 102:1274–1283, https://dx.doi.org/10.1016/j.bpj.2011.12.059

Pallavicini, C., A. Monastra, N.G. Bardeci, D. Wetzler, V. Levi, and L. Bruno. 2017. Characterization of microtubule buckling in living cells. Eur Biophys J, https://dx.doi.org/10.1007/s00249-017-1207-9

Pham, H.T.M. 2004. {PE}-{CVD} {S}ilicon {C}arbide - a structured material for surface micromachined devices. Doctoral Thesis, https://dx.doi.org/uuid:91e396a8-5857-4e93-ba27-8a82d0d9f2b7

Piedra, F.A., T. Kim, E.S. Garza, E.A. Geyer, A. Burns, X. Ye, and L.M. Rice. 2016. {GDP}-to-{GTP} exchange on the microtubule end can contribute to the frequency of catastrophe. Mol Biol Cell. 27:3515–3525, https://dx.doi.org/10.1091/mbc.E16-03-0199

Preciado Lopez, M., F. Huber, I. Grigoriev, M.O. Steinmetz, A. Akhmanova, G.H. Koenderink, and M. Dogterom. 2014. Actin-microtubule coordination at growing microtubule ends. Nat Commun. 5:4778, https://dx.doi.org/10.1038/ncomms5778

Rickman, J., C. Duellberg, N.I. Cade, L.D. Griffin, and T. Surrey. 2017. Steady-state {EB} cap size fluctuations are determined by stochastic microtubule growth and maturation. Proc Natl Acad Sci U S A. 114:3427–3432, https://dx.doi.org/10.1073/pnas.1620274114

Roth, D., B.P. Fitton, N.P. Chmel, N. Wasiluk, and A. Straube. 2018. Spatial positioning of {EB} family proteins at microtubule tips involves distinct nucleotide-dependent binding properties. J Cell Sci. 132, https://dx.doi.org/10.1242/jcs.219550

Schek, H.T., 3rd, M.K. Gardner, J. Cheng, D.J. Odde, and A.J. Hunt. 2007. Microtubule assembly dynamics at the nanoscale. Curr Biol. 17:1445–1455, https://dx.doi.org/10.1016/j.cub.2007.07.011

Schindelin, J., I. Arganda-Carreras, E. Frise, V. Kaynig, M. Longair, T. Pietzsch, S. Preibisch, C. Rueden, S. Saalfeld, B. Schmid, J.Y. Tinevez, D.J. White, V. Hartenstein, K. Eliceiri, P. Tomancak, and A. Cardona. 2012. Fiji: an open-source platform for biological-image analysis. Nat Methods. 9:676–682, https://dx.doi.org/10.1038/nmeth.2019

Seetapun, D., B.T. Castle, A.J. McIntyre, P.T. Tran, and D.J. Odde. 2012. Estimating the microtubule {GTP} cap size in vivo. Curr Biol. 22:1681–1687, https://dx.doi.org/10.1016/j.cub.2012.06.068

Sirajuddin, M., L.M. Rice, and R.D. Vale. 2014. Regulation of microtubule motors by tubulin isotypes and post-translational modifications. Nat Cell Biol. 16:335–344, https://dx.doi.org/10.1038/ncb2920

Taberner, N., G. Weber, C. You, R. Dries, J. Piehler, and M. Dogterom. 2014. Reconstituting functional microtubule-barrier interactions. Methods Cell Biol. 120:69–90, https://dx.doi.org/10.1016/B978-0-12-417136-7.00005-7

Tilney, L.G., J. Bryan, D.J. Bush, K. Fujiwara, M.S. Mooseker, D.B. Murphy, and D.H. Snyder. 1973. Microtubules: evidence for 13 protofilaments. J Cell Biol. 59:267–275, https://dx.doi.org/10.1083/jcb.59.2.267

Tischer, C., D. Brunner, and M. Dogterom. 2009. Force- and kinesin-8-dependent effects in the spatial regulation of fission yeast microtubule dynamics. Mol Syst Biol. 5:250, https://dx.doi.org/10.1038/msb.2009.5

Valiyakath, J., and M. Gopalakrishnan. 2018. Polymerisation force of a rigid filament bundle: diffusive interaction leads to sublinear force-number scaling. Sci Rep. 8:2526, https://dx.doi.org/10.1038/s41598-018-20259-7

VanBuren, V., L. Cassimeris, and D.J. Odde. 2005. Mechanochemical model of microtubule structure and self-assembly kinetics. Biophys J. 89:2911–2926, https://dx.doi.org/10.1529/biophysj.105.060913

VanBuren, V., D.J. Odde, and L. Cassimeris. 2002. Estimates of lateral and longitudinal bond energies within the microtubule lattice. Proc Natl Acad Sci U S A. 99:6035–6040, https://dx.doi.org/10.1073/pnas.092504999

Vleugel, M., M. Kok, and M. Dogterom. 2016. Understanding force-generating microtubule systems through in vitro reconstitution. Cell Adh Migr:475–494, https://dx.doi.org/10.1080/19336918.2016.1241923

von Loeffelholz, O., N.A. Venables, D.R. Drummond, M. Katsuki, R. Cross, and C.A. Moores. 2017. Nucleotide- and {M}al3-dependent changes in fission yeast microtubules suggest a structural plasticity view of dynamics. Nat Commun. 8:2110, https://dx.doi.org/10.1038/s41467-017-02241-5

Walker, R.A., N.K. Pryer, and E.D. Salmon. 1991. Dilution of individual microtubules observed in real time in vitro: evidence that cap size is small and independent of elongation rate. J Cell Biol. 114:73–81, https://dx.doi.org/10.1083/jcb.114.1.73

Waterman-Storer, C.M., J. Gregory, S.F. Parsons, and E.D. Salmon. 1995. Membrane/microtubule tip attachment complexes ({TAC}s) allow the assembly dynamics of plus ends to push and pull membranes into tubulovesicular networks in interphase {X}enopus egg extracts. J Cell Biol. 130:1161–1169, https://dx.doi.org/10.1083/jcb.130.5.1161

Zakharov, P., N. Gudimchuk, V. Voevodin, A. Tikhonravov, F.I. Ataullakhanov, and E.L. Grishchuk. 2015. Molecular and Mechanical Causes of Microtubule Catastrophe and Aging. Biophys J. 109:2574–2591, https://dx.doi.org/10.1016/j.bpj.2015.10.048

Zhang, R., G.M. Alushin, A. Brown, and E. Nogales. 2015. Mechanistic Origin of Microtubule Dynamic Instability and Its Modulation by {EB} Proteins. Cell. 162:849–859, https://dx.doi.org/10.1016/j.cell.2015.07.012

