## Supplemental material for "EB3-informed dynamics of the microtubule stabilizing cap during stalled growth"

Maurits Kok[a], Florian Huber[a,b,c], Svenja-Marei Kalisch[a], and Marileen Dogterom[a]*

[a] *Department of Bionanoscience, Kavli Institute of Nanoscience, Faculty of Applied Sciences, Delft Institute of Technology, van der Maasweg 9, 2629 HZ Delft, The Netherlands*

[b] *Netherlands eScience Center, Science Park 140, 1098XG Amsterdam, The Netherlands*

[c] *present affiliation: Center for Digitalization and Digitality, HSD – University of Applied Sciences, Münsterstraße 156, 40476 Düsseldorf, Germany*

**Key words**

microtubules, EB3, dynamic instability, *in vitro* reconstitution, 1D model, microfabricated barriers, Monte Carlo simulation

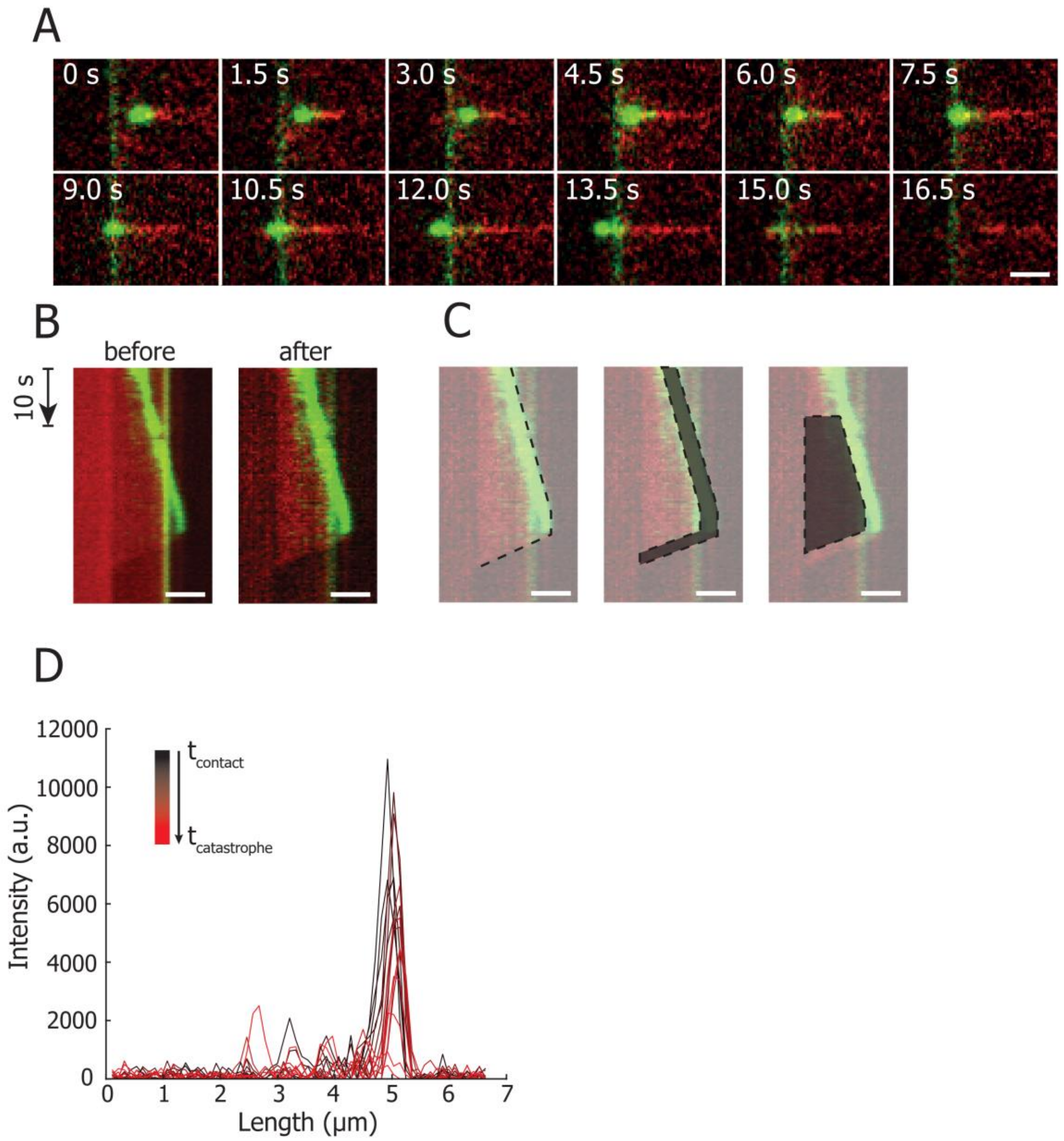


**Supplemental figure 1**

**A)** Montage of a microtubule-barrier stalling event in the presence of 15µM tubulin and 20nM EB3. During barrier contact microtubule polymerization halts, the EB3 signal at the microtubule plus end diminishes as GTP hydrolysis progresses, followed by a catastrophe. Scale bar denotes 5 µm.

**B)** In order to increase the signal to noise and remove light scattering at the edge of the overhang, a minimum projection of the image stack is subtracted from each pixel. The kymographs show the data before and after correction of the event in (A).

**C)** Determination of the position of the microtubule tip is obtained by manual tracking (left). Due to the loss of the EB signal during stalling and insufficient signal-to-noise of the microtubule signal, automatic tracking was not possible. The intensity of the EB3 comet is obtained from a region surrounding the manual trace with a width of 10 pixels or 1,1 μm (middle). The mean EB3 signal at the microtubule tip is corrected by subtracting the mean lattice intensity (right).

**D)** EB3 signal during microtubule-barrier contact of the event in **(A)**. The total contact duration is 3.75 seconds.

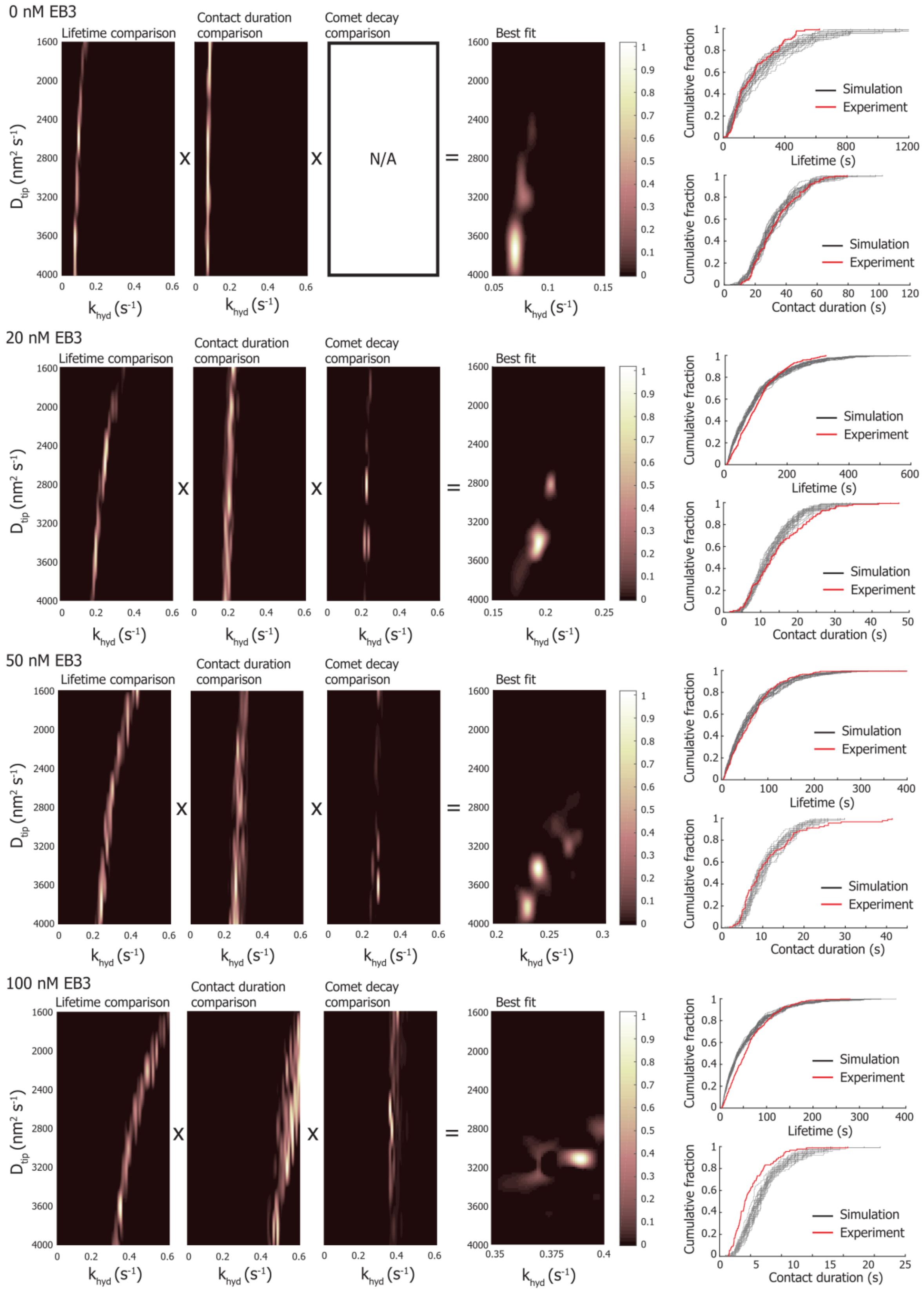


### Supplemental figure 2

In order to compare the distributions of the experimental and simulated microtubule lifetimes and stalling durations, we perform a Kolmogorov-Smirnov test. By simulating 500 events of

$D_{tip}$ and $k_{hyd}$ combinations for 0, 20, 50, and 100 nM EB3, we obtain (normalized) heatmaps of similarity for microtubule lifetime distributions and stalling duration distributions. The experimental and simulated decay rates were evaluated by their absolute difference. To determine the parameter set best capturing all observations, we calculate the product of the three comparisons. 25 bootstrapped simulated distributions based on the best fitting parameter set are plotted together with the experimental distribution.

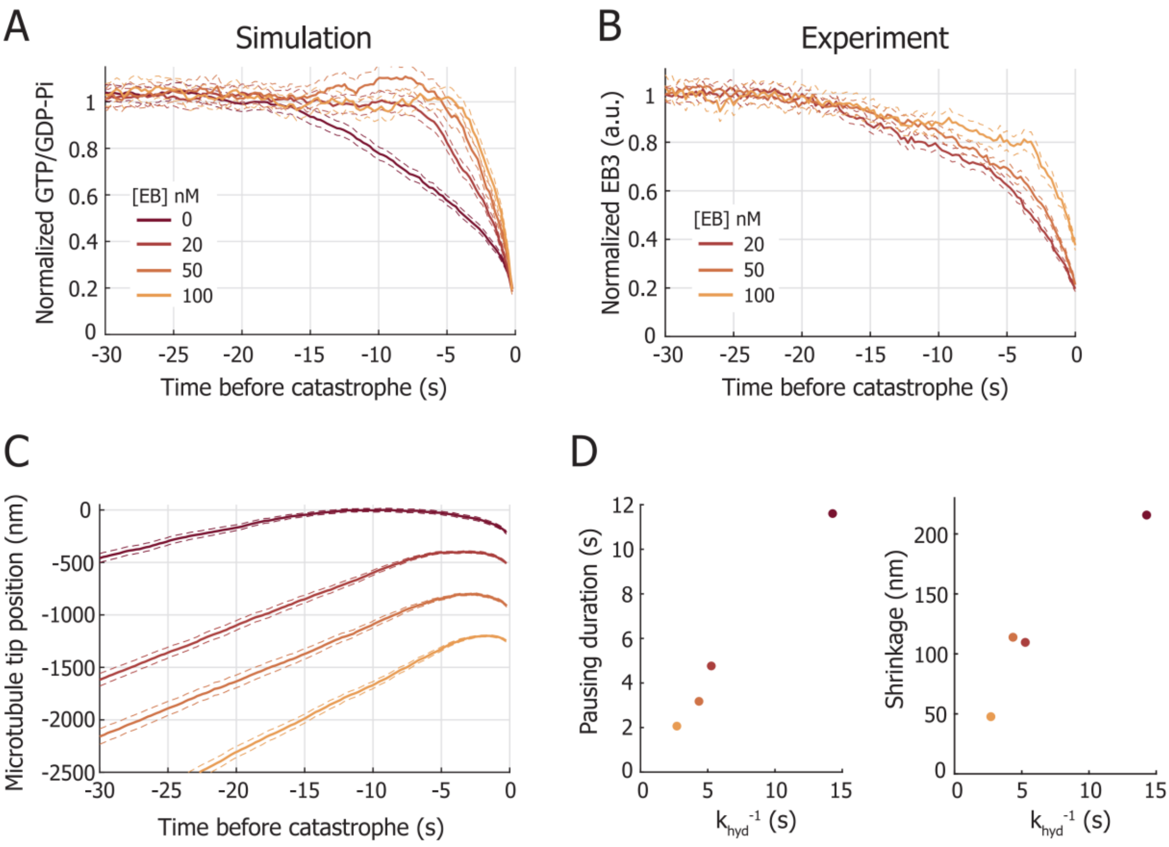


**Supplemental figure 3**

- **A)** Mean normalized GTP/GDP-Pi trace of 1000 simulated stalling events aligned on the moment of catastrophe (mean ± SE). The simulation parameters for 0, 20, 50, and 100 nM EB3 are found in Fig. 5C.
- **B)** Mean experimental EB3 intensity traces during microtubule stalling aligned on the moment of catastrophe (mean ± SE). The intensity is normalized with the mean of the steady-state intensity prior to barrier contact. Number of stalling events analysed: 20 nM, n = 151, 50 nM, n = 104, and 100 nM EB3, n = 92.
- **C)** Simulated position of the microtubule tip during microtubule stalling prior to catastrophe for 0, 20, 50, and 100 nM EB3 (mean ± SE).

**D)** Duration of pausing **(left)** and microtubule shrinkage **(right)** before the onset of catastrophe after barrier contact based on Fig. S3C. Microtubule pausing is defined as the time between the moment that the growth velocity is reduced to 10% of the steady-state value and the moment of catastrophe. Both the pausing duration and the shrinkage before the onset of catastrophe depend on the mean duration of hydrolysis.

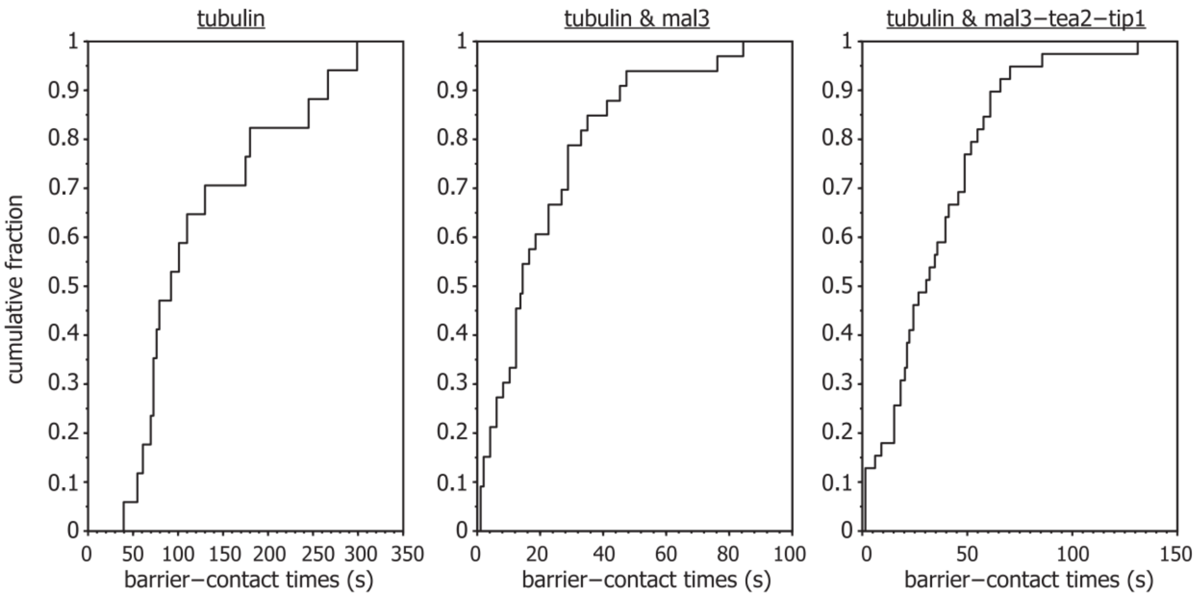


**Supplemental figure 4**

Previously unpublished data of the barrier contact duration using the barrier design from (Kalisch et al., 2011). The barriers were composed of SiO with a height of approximately 1.5 µm and without an overhang. Microtubules were nucleated from GMPCPP-seeds towards the barriers in the presence of 15 µM tubulin alone, and with addition of 200 nM Mal3-Alexa488, and with the combined addition of 200 nM Mal3-Alexa488, 8 nM Tea2, and 50 nM Tip1.

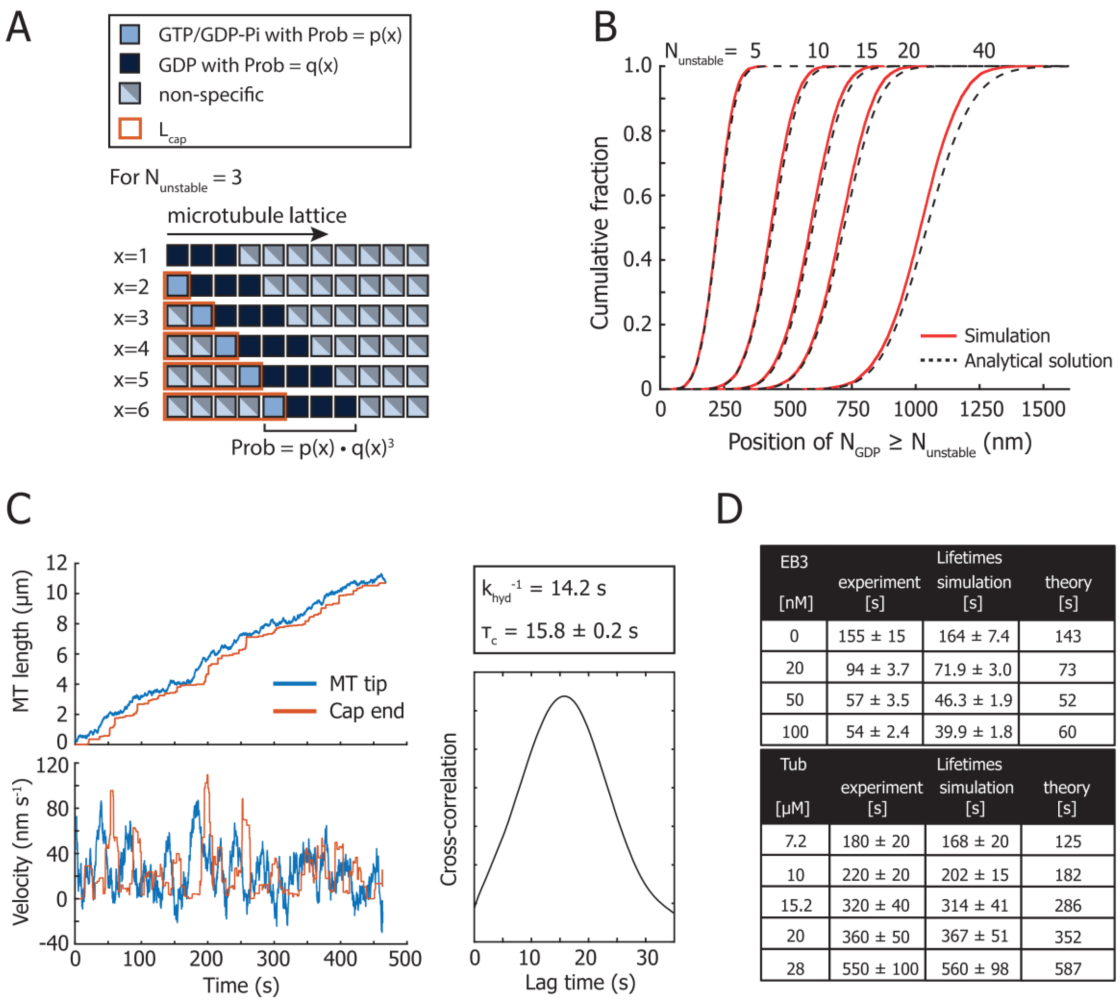


| EB3 [nM] | Lifetimes experiment [s] | Lifetimes simulation [s] | Lifetimes theory [s] |
|---|---|---|---|
| 0 | 155 ± 15 | 164 ± 7.4 | 143 |
| 20 | 94 ± 3.7 | 71.9 ± 3.0 | 73 |
| 50 | 57 ± 3.5 | 46.3 ± 1.9 | 52 |
| 100 | 54 ± 2.4 | 39.9 ± 1.8 | 60 |

| Tub [μM] | Lifetimes experiment [s] | Lifetimes simulation [s] | Lifetimes theory [s] |
|---|---|---|---|
| 7.2 | 180 ± 20 | 168 ± 20 | 125 |
| 10 | 220 ± 20 | 202 ± 15 | 182 |
| 15.2 | 320 ± 40 | 314 ± 41 | 286 |
| 20 | 360 ± 50 | 367 ± 51 | 352 |
| 28 | 550 ± 100 | 560 ± 98 | 587 |

**Supplemental figure 5**

**A)** Schematic of calculating the probability of finding a sequence of $N \geq N_{unstable}$ GDP subunits at a distance $x$ from the tip. The microtubule lattice is treated as a series of independent Bernoulli trials with probability $p(x)$ of finding a GTP/GDP-Pi subunit and probability $q(x) = 1 - p(x)$ of finding a GDP subunit. The probability of finding a sequence of 3 GDP subunits at position $x$ is $p(x)q(x)^3$.

**B)** The location of $N_{GDP} \geq N_{unstable}$ in the 1D microtubule lattice follows a Gaussian cumulative density distribution. The analytical solution holds well for the entire range of $N_{unstable}$ values explored in the simulations.

**C)** Correlation between the simulated growth fluctuations and the position of the cap end, as defined by $N_{unstable}$. Example of a simulated microtubule growth event showing the position of the microtubule tip and cap end, based on the parameters for 0 nM EB in Fig 5C. (**left, top**). The velocity of the cap end is correlated with the growth fluctuations of the microtubule tip (**left, bottom**). The cross-correlation between the tip and cap end velocity has a characteristic delay, which is approximately equal to ${k_{hyd}}^{-1}$.

**D)** Table with the microtubule lifetimes (median ± SEM) based on the EB-dependent dataset in Fig. 5C (**top**) and on the dataset in Fig. 7B (**bottom**) (Janson et al., 2003). We find that the experimental, simulated, and theoretical lifetimes are in good agreement.

A

| Experiment | | | |
|---|---|---|---|
| EB3 [nM] | Lifetime [s] | shape | rate [$s^{-1}$] |
| 0 nM | 155 ± 15 | 1.9 ± 0.23 | 0.0095 ± 0.001 |
| 20 nM | 94 ± 3.7 | 1.9 ± 0.12 | 0.018 ± 0.001 |
| 50 nM | 57 ± 3.5 | 1.6 ± 0.12 | 0.024 ± 0.002 |
| 100 nM | 54 ± 2.4 | 1.7 ± 0.10 | 0.027 ± 0.002 |

B

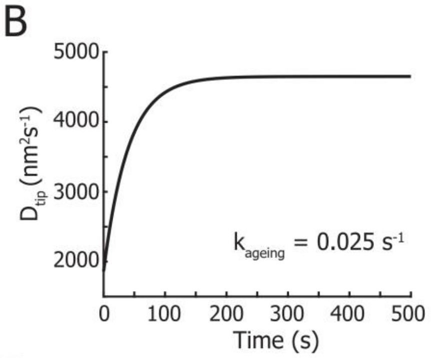


C

| Simulation | | | |
|---|---|---|---|
| EB3 [nM] | $k_{ageing}$ [$s^{-1}$] | shape | rate [$s^{-1}$] |
| 0 nM | 0.025 | 1.7 ± 0.19 | 0.0085 ± 0.001 |
| 20 nM | 0.025 | 1.9 ± 0.22 | 0.019 ± 0.003 |
| 50 nM | 0.025 | 2.0 ± 0.24 | 0.029 ± 0.004 |
| 100 nM | 0.025 | 2.1 ± 0.24 | 0.027 ± 0.004 |

D

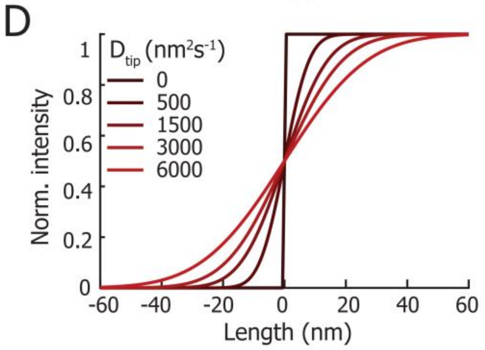


**Supplemental figure 6**

**A)** Table with experimental microtubule ageing parameters. The shape and rate parameters were obtained by fitting the lifetime distributions for freely growing microtubules with a Gamma distribution (mean ± 95% CI).

**B)** Time-dependent tip fluctuations are sufficient to introduce microtubule ageing. The tip fluctuations increase according to $D_{tip}(t) \propto 1 - e^{-0.025t}$, with $D_{tip}(0) = \frac{1}{2} D_{tip}$ and $D_{tip}(\infty) = \frac{5}{4} D_{tip}$.

**C)** Table with simulated microtubule ageing parameters. The shape and rate parameters were obtained by fitting the lifetime distributions for freely growing microtubules with a Gamma distribution (mean ± 95% CI).

**D)** Theoretical shape of an experimentally measured microtubule tip. Fluctuations of the microtubule could generate a blurred during frame acquisition with TIRF. However, the calculated blurring is on the order of 20 nm, too small to be detected by TIRF microscopy.

**Supplemental movies**

**Movie S1, related to figure 1. Microtubules polymerizing towards the barrier in the presence of tubulin alone.**

Microtubules are growing towards the microfabricated barriers in the presence of 15 µM HiLyte488-tubulin. The GMPCPP seeds are labelled in magenta and the tubulin in green. Images were collected with TIRF microscopy at a 500 ms interval. Video is sped up 120 times. Time is shown in the format min:sec.

**Movie S2, related to figure 1. Microtubules polymerizing towards the barrier in the presence of GFP-EB3.**

Microtubules are growing towards the microfabricated barriers in the presence of 15 µM rhodamine-tubulin (magenta) and 20 nM GFP-EB3 (green). Images were collected with TIRF microscopy at a 250 ms interval. Video is sped up 240 times. Time is shown in the format min:sec.
